# Operon formation by insertion sequence IS*3* in *Escherichia coli*

**DOI:** 10.1101/2021.11.02.466885

**Authors:** Yuki Kanai, Saburo Tsuru, Chikara Furusawa

## Abstract

Operons are a hallmark of the genomic and regulatory architecture of prokaryotes. However, the mechanism by which two genes placed far apart gradually come close and form operons remains to be elucidated. Here, we propose a new model of the origin of operons: Mobile genetic elements called insertion sequences can facilitate the formation of operons by consecutive insertion-deletion-excision reactions. This mechanism barely leaves traces of insertion sequences and is difficult to detect in evolution in nature. We performed, to the best of our knowledge, the first experimental demonstration of operon formation, as a proof of concept. The insertion sequence IS*3* and the insertion sequence excision enhancer are genes found in a broad range of bacterial species. We introduced these genes into insertion sequence-less *Escherichia coli* and found that, supporting our hypothesis, the activity of the two genes altered the expression of genes surrounding IS*3*, closed a 2.7 kilobase pair gap between a pair of genes, and formed new operons. This study shows how insertion sequences can facilitate the rapid formation of operons through locally increasing the structural mutation rates and highlights how coevolution with mobile elements may shape the organization of prokaryotic genomes and gene regulation.

## INTRODUCTION

Operons are clusters of genes under the control of the same promoter sequences and are a hallmark of prokaryotic genome structure (1, 2) and gene regulation (3). A significant proportion of genes of the prokaryotic genome are organized into operons (4), and it is widely accepted that operons are beneficial for multiple reasons, including for better coregulation of genes (5–8).

On the other hand, the mechanism by which operons form is poorly understood (9). As understanding operon formation is fundamental to our understanding of the evolution of prokaryotes, many models for operon formation have been proposed in the last 60 years (7, 9-12). However, none of the proposed mechanisms of operon formation have been corroborated by experimental evidence, limiting our understanding of the types of mutations that drive them.

In principle, operons form by rearrangements, including insertions, deletions (9) (Figure 1.A), and duplications (10), but these mechanisms alone seem to be insufficient to explain the prevalence of operons in prokaryotic genomes. For instance, duplication may explain the evolution of some operons (10), but genes in operons do not necessarily have similar structures (7). Moreover, while pairs of genes can culminate into operons by random rearrangements, they are more likely to end up apart (7).

**Figure 1.**
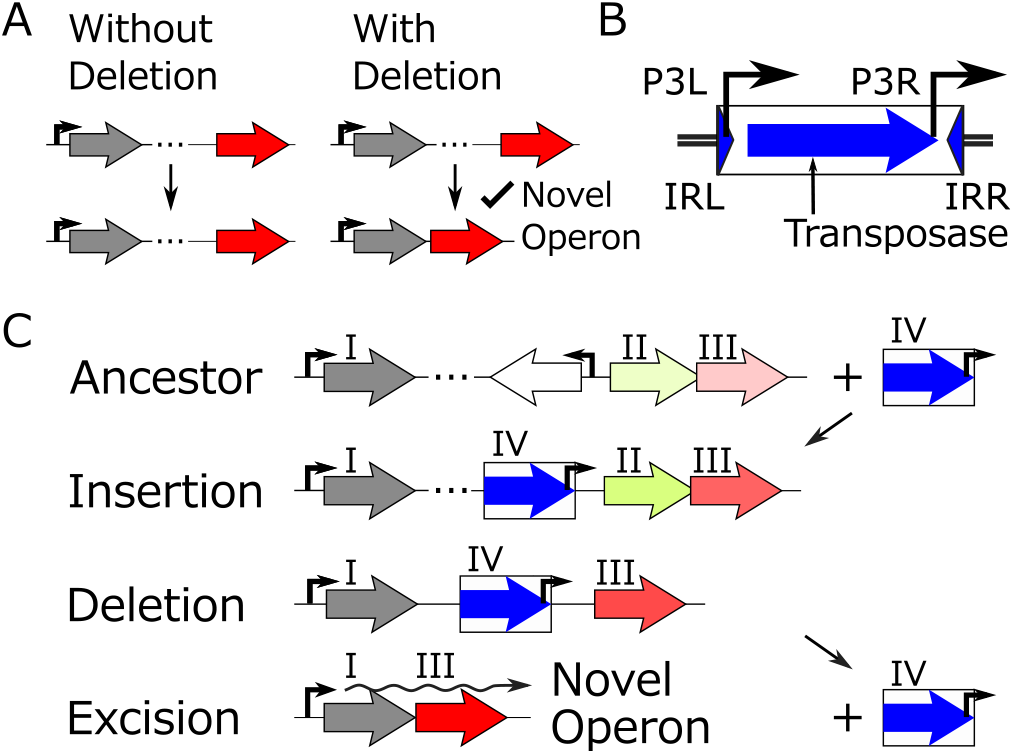
Insertion sequences (ISs) can facilitate the formation of novel operons through insertion-deletion-excision reactions (IDE model). **(A)** Deletion facilitates the formation of operons. **(B)** IS*3* has two promoters, P3L and P3R, in the same direction. **(C)** An example of how ISs (IV) can facilitate the formation of operons based on the IDE model. The block arrows indicate coding sequences, and the intensity of green (II) and red (III) colors indicates the gene expression level. In an environment where the genes indicated in red (III) and gray (I) are essential and that in green (II) is nonessential, the reactions from the top to bottom leads to the genes in gray and red to form a novel operon.

Thus, selection for gene coregulation is thought to be crucial after operon formation by the aforementioned mechanisms to account for the prevalence of operons (5, 9, 13, 14). However, while selection for coregulation can facilitate the maintenance and sophistication of operons (9), it does not facilitate their formation (7). This is because bringing two genes closer becomes adaptive only when the genes were initially clustered close enough (7). The ‘selfish operon’ model proposed that instead of relying on rare random rearrangements to cluster genes, operons form gradually through small intermediate steps (7). Unfortunately, this model has not found general acceptance as it fails to explain how essential genes likely form operons (5, 15). Operon formation likely proceeds through a mechanism that gradually ties two far apart genes (7); however, a plausible mechanism remains to be elucidated.

As genes in operons are clustered together, mechanisms that cluster functionally related genes may also facilitate the formation of operons (13). Essential genes in prokaryotic genomes seem to cluster because the genomes tend to lose tandem sequences (‘persistence model’ (12)); Clustering of essential genes increases the robustness of genomes against tandem deletions by making large deletions less detrimental (16). Recently, a laboratory evolution of *E. coli* revealed that the major cause of tandem deletions may be the activity of insertion sequences (IS, Figure 1.B) (17). This could mean that the tandem deletions by IS cluster important genes in prokaryotic genomes and may also make those genomes operon-rich.

ISs are a major cause of genetic rearrangements (17, 18) and therefore may as well facilitate operons to form. A recent study, based on the presence of ISs within ancient operons, inferred that ISs may form operons by facilitating rearrangements (19). However, no study seems to have associated the tandem deletion of surrounding genes by IS with the formation of operons.

Although ISs are known to delete neighboring sequences since at least the 1970s (20–22), deletions by IS as a mechanism that promotes the formation of operons have been undervalued for the following three main reasons: (i) IS-rich genomes seem to have more operons disrupted compared to other genomes (19, 23). Therefore, ISs are often associated with the disruption rather than the formation of operons. (ii) IS excision is required for an operon to form by tandem deletion (Figure 1.C). However, IS excision would result in a double-strand break in the genome, which may lead to genomic degradation unless fixed by homologous recombination (24). (iii) Some of the major types of ISs rarely excise themselves (25).

The following findings have made these arguments less critical: (i) IS-rich genomes can be rich in operons (26), suggesting that IS-mediated rearrangements may not only destroy but also form operons (9). (ii) Many bacterial species have end-joining mechanisms (27). (iii) Many ISs are known to excise themselves (28, 29). Moreover, the excision of one of the major types of IS, IS*3*, that rarely excise themselves, is significantly promoted by an enzyme called IS excision enhancer (IEE) found in various bacterial species (25).

We hypothesized that, considering that IS*3* is known to rapidly create deletions of various lengths adjacent to it (21, 30), the interaction of IS*3* and IEE may promote the rapid formation of operons by IS*3* deleting sequences intervening two genes and IEE excising IS*3*.

In this study, we assess if deletion by IS can promote the formation of operons. First, we propose a model of operon formation based on deletions by ISs, that does not rely on the selection for coregulation. We also suggest a mechanism by which an IS can facilitate the formation of operons through intermediate steps, gradually bringing closer two genes placed far apart. According to this model, ISs do not leave apparent traces after forming operons. This makes it difficult to demonstrate the model using comparative genomics of extant genomes. Therefore, as a proof of concept, we engineered *E. coli* in a state that mimics an intermediate step of the model by inserting a copy of IS*3* between two conditionally essential genes. To detect the diversity of structural mutations during the formation of operons, fluorescent reporters were placed around the copy of IS*3*. Selecting the cells cultured overnight based on their fluorescence, we examined how the concerted activity of IS*3* and IEE affects the expression of surrounding genes, leading to the formation of novel operons. We believe this study is the first to provide experimental evidence of a plausible mechanism of operon formation in prokaryotes.

## MATERIALS AND METHODS

### Reagents

#### For cell culture, we used

LB medium (Difco LB Broth, Miller, BD, USA, 244620), kanamycin (Kanamycin Sulfate, Wako, Japan, 115-00342), streptomycin (Streptomycin Sulfate, Wako, Japan, 190-14342), ampicillin (Ampicillin Sodium, Wako, Japan, 012-23303), anhydrotetracycline hydrochloride (aTc) (Anhydrotetracycline hydrochloride, Sigma-Aldrich, USA, 37919), arabinose (L(+)-Arabinose, Wako, Japan, 010-04582), IPTG (Isopropyl-*β*-D(-)-thiogalactopyranoside [IPTG], Wako, Japan, 190-14342), agar (Agar, Powder, Wako, Japan, 010-08725).

#### For handling nucleic acids and DNA sequencing, we used

KOD One PCR Master Mix Blue (Toyobo, Japan, KMM-201), Rapid Barcoding Kit (Oxford Nanopore Technologies, UK, SQK-RBK004), DpnI (Takara Bio, Japan, 1235-A), PureLink RNA Mini Kit (Thermo Fisher Scientific, USA, 12183018A), PureLink DNase Set (Thermo Fisher Scientific, USA, 12185010), PrimeScript High Fidelity RT-PCR Kit (Takara Bio, Japan, R022A), In-Fusion Snap Assembly Master Mix (Takara Bio, Japan, 638948), FastGene Plasmid Mini Kit (NIPPON Genetics, Japan, FG-90502).

#### We used the following instruments

Infinite F200 multimode plate reader (Tecan, Switzerland), Flongle Flow Cell R9.4.1 (Oxford Nanopore Technologies, UK, FLO-FLG001), MinION Mk1B (Oxford Nanopore Technologies, UK, MIN-101B), FACSAria III (BD, USA), MiniAmp Plus (Thermo Fisher Scientific, USA), blue/green LED transilluminator (NIPPON Genetics, Japan, LB-16BG).

The primers we used are listed in Supplementary Table 1.

### Biological resources

#### We used the following strains

*E. coli* MG1655 (the origin of IS*3* transposase used in this study), *E. coli* MDS42 (the parent strain used for laboratory evolution, (31)), *E. coli* HST08 (used to test coregulation of genes in newly-formed operons and for vector construction; *E. coli* HST08 Premium Competent Cells, Takara Bio, Japan, 9128).

We used several plasmids in this study. pCDSSara-L18 (32) was used as the backbone of pYK-1S3 and as a –*iee* control. The Tn5 transposase gene of pRC2117 (33) was inserted instead of IS*3* transposase as a –*tpn* control. The following plasmids were constructed for this study: pYK-1N5 (the fluorescence reporter plasmid), pYK-1S3 (the IEE expression plasmid), pYK-1Q2 (pYK-1N5 with Tn5 transposase gene within a pair of IS*3* inverted repeats), pYK-1S0 (plasmid constructed based on pYK-1N5 but with a P*_tac_* cassette).

### Strains and plasmids

To identify the effect of IS*3* on the formation of operons, *E. coli* MDS42, a derivative of the wild-type K-12 MG1655, was used as it is absent of all mobile elements, including IS*3* (31). IS*3* and IEE were supplied via plasmids pYK-1N5 and pYK-1S3, respectively. All plasmids were constructed using the In-Fusion cloning kit.

The IS*3* supplying vector (pYK-1N5, Figure 2.A) was designed to mimic a state after the ‘insertion’ step of the model that we propose in this study (Figure 1.C). pYK-1N5 is a low copy-number plasmid (pSC101 origin) with a kanamycin resistance gene (*kanR*), an engineered IS*3* sequence, and red (mScarlet-I (34)) and yellow (Venus YFP (35)) fluorescent protein genes (*yfp, rfp*). To demonstrate our proposed model, IS*3* was engineered to have between its two inverted repeats (IR) the IS*3* transposase gene (*tpn*) downstream of an inducible promoter (based on P_LtetO-1_ (36, 37)) and a synthetic promoter P_J23105_ (a Biobrick promoter). The P_LtetO-1_ derivative is repressed by the product of *tetR* on the same plasmid, and therefore, transcription is inducible by the addition of aTc to the growth medium. P_J23105_ was inserted to mimic the outward-facing promoter P3R of wildtype IS*3* because, according to our model, it facilitates the formation of operons. We chose P_J23105_ because its strength was ideal for demonstrating using flow cytometry (FCM) the influence of IS*3* activity on the expression of surrounding genes. The IS*3* used in this study was designed so that the two promoters of IS*3* were divergent, with the left IR (IRL) downstream of the transposase gene. This design was chosen to avoid induction by aTc to increase the transcription of the fluorescent protein, to avoid potential promoter activity from IRL to interfere with P_LtetO-1_, and to prevent the plasmid from being degraded before evolution by inhibiting the activity of the transposase gene by antisense transcription from the *kanR* gene. In addition, following a previous study (21), the two open reading frames constituting the IS*3* transposase gene were fused by a single base insertion in the A_4_G motif to increase the activity of the transposase.

**Figure 2.**
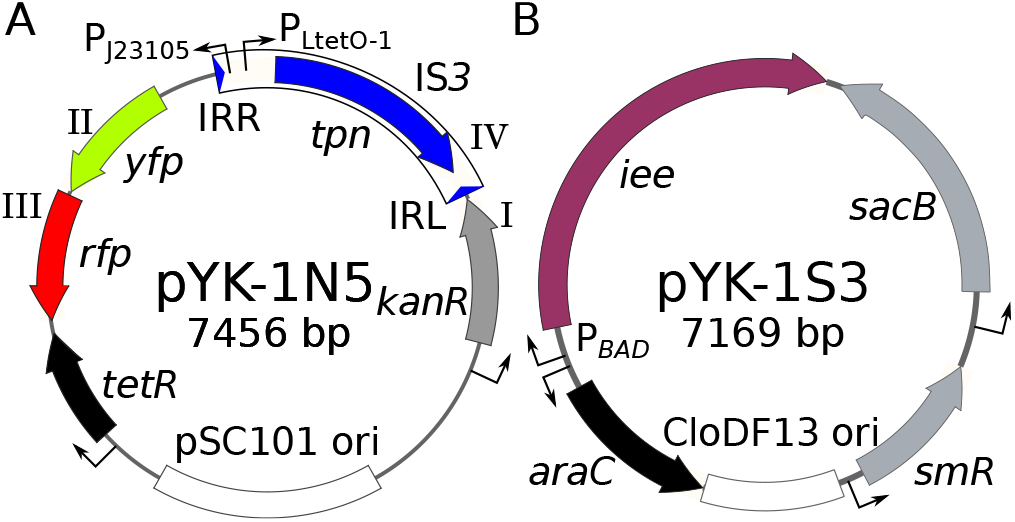
**(A)** Design of the fluorescent reporter plasmid used to detect the formation of operons. The plasmid was designed to mimic the ‘insertion’ step of operon formation shown in Figure 1.C. See *strains and plasmids* for details. **(B)** Design of the insertion sequence excision enhancer (IEE) expression plasmid. Tetracycline repressor gene, *tetR*; origin of replication, ori; arabinose operon regulator gene, *araC*; streptomycin resistance gene, *smR*. The genes of pYK-1N5 with roman numerals correspond to the genes in Figure 1.C with the same roman numerals.

The IEE expression vector (pYK-1S3, Figure 2.B) was constructed based on pCDSSara-L18 (32) by replacing the arabinose-inducible ribosomal protein gene with *iee* of *E. coli* O157:H7 strain Sakai (NCBI Gene ID: 912859, ECs_1305). This plasmid had a streptomycin resistance gene (*smR*) as a selection marker.

Derivatives of the two plasmid vectors without the IS*3* and IEE genes were used as negative controls. A hyper-active mutant of Tn*5* transposase gene (33) and ribosomal subunit L18 (32) were placed instead of IS*3* transposase and IEE genes, respectively.

To introduce IS*3* and IEE, the ancestor strains for evolution were prepared by two rounds of electrotransformation. First, pYK-1S3 (or its –*iee* derivative) was introduced by transformation, and the transformed cells were selected on LB agar plates containing streptomycin. Mixtures of electrocompetent cells were then prepared from the selected colonies. Finally, pYK-1N5 (or its –*tpn* derivative) was introduced by transformation into the cells and the transformed cells were grown on LB agar plates containing streptomycin and kanamycin.

### Culture conditions

Cells were cultured aerobically in 200 μL volume of LB medium in 96-well plates (Greiner Bio-One, Cellstar, 655180). First, the medium was filtered through a 0.2 μm membrane filter (Thermofisher Scientific, 567-0020) to remove debris that may interfere with FCM measurements. The culture medium was supplemented with 50 μg/mL kanamycin and 100 μg/mL streptomycin where indicated, to select only cells with both pairs of plasmids. To induce transcription of transposase and IEE genes, 50 nM of aTc and 0.1 % (w/v) of arabinose, respectively, were added to the media where indicated.

Growth rates were measured in Infinite F200 multimode plate reader at 37 °C overnight. The plate was shaken linearly and then orbitally for three minutes each, and OD_600_ was measured every ten minutes to determine the growth rate.

To obtain colonies, cells were plated on 1.5 % agar plates supplemented with LB and appropriate antibiotics.

Cells were temporarily stored at 4 °C as colonies grown on LB agar plates or as mixtures in filtered phosphate-buffered saline (PBS).

### Flow cytometry and selection of cells by fluorescence-activated cell sorting (FACS)

For the evolution of the cells, four colonies of each of the four genotypes (± *tpn* and ± *iee*) were picked with toothpicks from LB agar plates, diluted 100-fold, and cultured overnight in 96-well plates with 200 μL LB containing streptomycin and kanamycin supplemented with aTc and arabinose. After overnight culture at 30 °C, the cells were diluted in ice-cold PBS and stored at 4 °C until FCM measurements. The singlecell fluorescence of these cells was measured by FCM using FACSAria III and FACS Diva software v.6.1.3. The analysis and visualization were based on single-cell fluorescence of cells with forward scatter measurements within a twofold range, including the most frequent values to compare fluorescence among cells in similar physiological states (38).

For FACS, four populations of cells with both *tpn* and *iee* and one population of cells with *tpn* but without *iee* were prepared as described above. After a brief measurement of the distribution of fluorescence by FCM using 10,000 cells, gates were manually set by first subsetting the measurements with the same forward scattered values as above and then manually drawing the gates according to the measurement, as shown in Figure 4.A (P1-3). The lines of the gates were made thicker for visualization using Inkscape. The same gates were used for all five rounds of FACS. The cells were collected in 500 μL of LB or PBS.

### Genotyping cells

To confirm that the cells were sorted as expected, 200–500 μL of the collected cells were cultured overnight in 5 mL of LB medium containing kanamycin and streptomycin, and 10 μL of cultured cells were plated on LB agar plates with kanamycin and streptomycin. The plates were incubated at 37 °C for two days and imaged under a transilluminator. The brightness and contrast were adjusted using ImageJ (National Institutes of Health, USA).

Sanger sequencing was used to genotype the cells in each FACS gate. Colonies on agar plates showing uniform fluorescence were picked, and colony PCR was performed using KOD One PCR Master Mix Blue. Template DNA was prepared by picking the colonies by toothpicks into Tris-HCl (pH8.5) with 0.1 % Triton X-100, boiling at 98 °C for five minutes, and spinning down briefly at 10,000 ×g. The supernatants were used as templates. Agarose gel electrophoresis was run for the amplified DNA, and fragments of typical lengths were chosen and sequenced by Sanger sequencing (GeneWiz). The same procedure, but with a different primer set, was followed to check that sequences were not deleted in plasmids of cells in the major dark fraction of FCM measurements in +*iee* +*tpn* cells.

To identify the diversity among the cells sorted by FACS, nanopore sequencing was performed. 400 μL out of 500 μL of PBS with 600 cells collected by a single round of FACS were transferred into 5 mL of LB medium containing kanamycin and cultured overnight at 37 °C. Streptomycin was not added to reduce the read mapping onto pYK-1S3 and increase read mapping onto pYK-1N5. Plasmid DNA was extracted with FastGene Plasmid Mini Kit (with optional washing), and the sample for sequencing was prepared by adding barcodes and adapters using Rapid Barcoding Kit following the manufacturer’s instructions. The samples were applied to Flongle Flow Cell R9.4.1 on MinION Mk1B and sequenced using MinKNOW (v21.06.0). The fast5 data was basecalled using ONT Guppy (v5.0.11) with the configuration set to the ‘superaccurate model.’ The reads were filtered according to the default quality threshold, and the barcode sequences were trimmed. We obtained 3,522 reads with 14,705,564 base pairs (bp) corresponding to a sequencing depth of approximately 2,000. The median read length and quality were 4,359 bp and 13.03, respectively. Reads longer than one kbp were filtered with Filtlong (v0.2.0, https://github.com/rrwick/Filtlong) and mapped onto a fasta file with two copies of the pYK-1N5 sequence placed in tandem using minimap2 (v2.17-r941 (39)) to avoid reads that were mapped across the origin being split. Deletions longer than 50 bp were detected and extracted from the CIGAR strings of BAM using Rsamtools (v2.6.0) and analyzed using a custom script. The 514 reads are plotted in Figure 4.F.

### Reverse transcription polymerase chain reaction (RT-PCR)

To assess if *kanR* and *rfp* formed an operon, RT-PCR was performed. Two colonies genotyped by Sanger sequencing were picked into LB medium containing streptomycin and kanamycin and cultured overnight at 30 °C. The cultured cells were diluted 50-fold into a fresh growth medium and cultured at 30 °C for two hours. The total RNA was extracted using the PureLink RNA Mini Kit. DNA was removed using DNAse during the extraction. Total RNA was amplified using PrimeScript High Fidelity RT-PCR Kit following the manufacturer’s two-step protocol. First, reverse transcription was performed. RNA with a final concentration of approximately 50 ng/μL was denatured at 65 °C for five minutes with either of the forward and reverse primers and, as a negative control of reverse transcription, with neither of the primers; reverse-transcriptase and buffers were added on ice; reverse transcription was performed at 42 °C for 30 minutes; the reverse transcriptase was denatured at 95 °C for five minutes; the RNA mixtures were stored at 4 °C.

Then, cDNA was amplified by PCR. The obtained cDNA was diluted ten-fold with a premix containing the pair of primers and the DNA polymerase; DNA was amplified by 20 cycles of PCR (MiniAmp Plus), with denaturation at 98 °C for ten seconds, followed by annealing at 57 °C for five seconds, and extension at 72 °C for three minutes. The PCR products were subject to gel electrophoresis, and the image was recorded under a transilluminator. The brightness and contrast of the image were adjusted using ImageJ.

### Testing the coregulation of *kanR* and *rfp* in newly-formed operons

To further confirm operon formation after evolution, the *kanR* promoter of pYK-1N5 and its evolved derivatives were replaced with a lactose inducible promoter P_*tac*_. DNA sequences, excluding the kanamycin promoter sequence, of the plasmids were amplified by PCR (KOD One PCR Master Mix Blue). Another sequence containing the ampicillin resistance gene, P*_tac_*, and *lacI* (Supplementary Figure 6) was amplified by PCR from plasmid pYK-1S0. After digestion of remnant plasmids with DpnI, the amplified sequences were fused with In-Fusion cloning following the manufacturer’s manual, and the DNA mixture was introduced to *E. coli* HST08 Premium Competent Cells. The sequence of the insert is provided in the *DNA Sequences* section of *Supplementary Data*.

Cells with the re-engineered plasmids were spread on LB agar plates with ampicillin. After checking the genotypes of the cells, their colonies were picked and transferred to 96-well plates with 200 μL of LB supplemented with ampicillin and 30 μM of IPTG. The cells were cultured overnight at 32 °C to prevent edge effects. After the cells reached the stationary phase, the cells were diluted in ice-cold PBS, and single-cell fluorescence was measured using FCM.

### Statistical Analyses

Type III ANOVA was performed using FCM measurements of 16 independent cell cultures consisting of four colonies for each of the four genotypes. Because there were three treatments, there were 12 degrees of freedom within groups.

For calculating the number of cells in the three gates used for FACS, the mean and 95 % confidence interval (CI) of log10 probability are presented (n=4), assuming the normality of the distributions.

To show that IPTG induction increased red fluorescence, three independent cell cultures were prepared for each condition and genotype, and the intensity of single-cell red fluorescence was measured with FCM. First, the FCM data was subsetted based on their forward scatter values as above. Using the values of median log_10_ red fluorescence of over 10,000 measurements per culture, we performed two-sided t-tests adjusted with the Bonferroni method.

### Data Availability/Sequence Data Resources

Data related to nanopore sequencing and its analysis, including the fast5 data, reference fasta, and BAM files, were deposited in NCBI Sequence Read Archive (BioProject PRJNA768397). The raw flow cytometry data are available in FlowRepository (FR-FCM-Z4LV). DNA sequences of pYK-1N5, pYK-1S3, and the P*_tac_* cassette are provided in the *DNA sequences* section of *Supplementary Data*.

### Data Availability/Novel Programs, Software, Algorithms

Not applicable.

### Web Sites/Data Base Referencing

As references, we used [NCBI Gene](https://www.ncbi.nlm.nih.gov/gene/912859) for the DNA sequence of *iee*, and the [Registry of Standard Biological Parts](http://parts.igem.org/) to find various Biobrick promoter sequences, including P_J23105_.

## RESULTS

### A model of operon formation based on the activity of insertion sequences

To explain the formation of operons in prokaryotes, we propose a model that relies on the activity of ISs and experimentally demonstrate the formation of an operon. ISs may be able to cluster two genes placed far apart into an operon by a sequential Insertion-Deletion-Excision reaction as follows (IDE model, Figure 1.C).

#### Ancestor

Initially, two genes beneficial in the same environment are placed apart, as is the case for the genes in gray and red (I, III) in Figure 1.C.

#### Insertion

Many ISs have promoters that facilitate the transcription of downstream genes (40, 41). For example, in addition to the promoter upstream of the transposase gene, IS*3* has a strong promoter facing outward (P3R, Figure 1.B (42)). These promoters help an IS (IV) to activate initially dormant genes (II, III) by transposing upstream of the genes (41).

#### Deletion

Many types of ISs are known to delete sequences adjacent to them (20–22). Deletion can continue until nonessential genes between the IS and beneficial genes are deleted. This results in bringing together two beneficial genes on either side of the IS closer.

#### Excision

Many ISs can excise themselves, especially in the presence of IEE (25). This excision of an IS between two beneficial genes can lead to them forming a novel operon.

Thus, with the tandem deletion caused by IS, selection for the preservation of two genes (I and III) can lead to the formation of operons.

### The activity of IS can gradually form an operon through intermediate adaptive steps

Mechanisms that gradually bring closer two genes seem to be more effective than rare rearrangements that directly cluster two genes to form an operon (7). Our model involves such a mechanism and can be broken down into small intermediate steps, all of which are adaptive (Figure 1.C). For example, let us assume that the stronger expression of genes indicated in gray (I) and red (III) is beneficial and that initially, the gene in gray is upregulated, but the gene in red lacks an effective promoter (ancestor). The insertion of an IS (IV) upstream of the gene in red would increase the fitness, as the gene would be transcribed from the promoters within the IS. Further activity of the IS can delete sequences around the IS, increasing the expression of the gene in red by bringing it closer to the inner promoters of the IS and gradually closing the space between the IS and the gene. Adaptive evolution can also bring the two genes closer together by increasing the robustness of cells against tandem deletions or activity of selfish elements (12, 43). Finally, the excision of IS would be most beneficial when cells with both gray and red genes are adjacent to the IS as it would bring the promoter of the gene in gray closer to the gene in red. The premature excision of IS is less likely to be fixed because the gene in red would lose active promoters encoded within the IS.

The IDE reactions locally accelerate rearrangements, and while they do not require the selection for better gene coregulation to form operons as long as active transcription of the two genes is maintained, adaptive evolution can facilitate operon formation by fixing intermediate steps towards forming operons (7).

### A fluorescence reporter system to detect deletions related to the formation of *kanR-rfp* operon

To experimentally demonstrate our model of operon formation, we designed a plasmid as a model of the prokaryotic genome after the ‘insertion’ step (pYK-1N5, Figure 2.A), which is common in both nature and laboratory settings (30, 41, 44). Specifically, the fluorescent reporter plasmid pYK-1N5 was designed such that the kanamycin resistance gene (*kanR*, I), a copy of IS*3* (IV), and the red (*rfp*, III) and yellow (*yfp*, II) fluorescent protein genes correspond to the genes in gray, the copy of IS, and the genes in green and red in Figure 1.C, respectively. The copy of IS*3* was inserted within a 2.7 kbp gap between *kanR* and *rfp*. For ease of the experiment, the wild-type IS*3* was modified such that its transposase gene (*tpn*) was inducible by aTc, and the promoter P_J23105_ activated *rfp* instead of the native promoter of IS*3*. We expected that the IS would delete surrounding sequences or excise itself, changing the pattern of fluorescence measured by FCM. Using FACS, we selected cells with bright red fluorescence while adding kanamycin to the growth medium, expecting the formation of *kanR-rfp* operons.

With the two fluorescent protein genes, various phenotypes were expected to be found after evolution. Deletions between IS*3* and *yfp* would bring P_J23105_ closer to *yfp*, causing both yellow and red fluorescence to increase. Excision of IS*3* would lead to the transcription of the two fluorescent protein genes from the *kanR* promoter instead of the weaker P_J23105_, also causing both yellow and red fluorescence to increase. In this case, we expected *kanR-yfp-rfp* operon, or in case of *yfp* deletion, *kanR-rfp* operon to form.

### The pattern of fluorescence observed under the presence of both IS*3* and IEE

As a potential genetic background that facilitates operon formation, we examined whether the combined activity of IS*3* and IEE in *E. coli* can accelerate operon formation. IS*3* rarely excises itself on its own but is frequently excised under the presence of IEE (25). The presence of IEE thus is expected to promote the ‘excision’ step of operon formation. To test this, an arabinose-inducible IEE expression vector (pYK-1S3, Figure 2.B), in which IEE was expressed from the promoter of *araBAD* operon (P_BAD_), was constructed. As negative controls, derivatives of pYK-1N5 and pYK-1S3 without *tpn* and *iee*, respectively, were also constructed. We transformed an IS-less strain of *E. coli* MDS42 (31) with pairs of plasmids with and without *tpn* and *iee*. We expected that with both *tpn* and *iee*, a large proportion of cells would show an increased level of yellow and red fluorescence by losing the copy of IS*3*. We also expected some cells to show an increase in only red fluorescence as IS*3* can disrupt *yfp*.

To test if the activity of IS and IEE creates the expected fluorescence patterns, cells transformed with the pairs of plasmids were cultured overnight with both *iee* and *tpn* fully induced (Figure 3, Supplementary Figure 1). We found that when both *tpn* and *iee* were present, a fraction of cells showed intense yellow and red fluorescence (Figure 3, white arrowhead). While these cells were also found in other conditions, they were most apparent when both *tpn* and *iee* were present and fully induced. To validate the importance of having both *tpn* and *iee*, Type III ANOVA was performed with the proportion of measurements with red fluorescence ten times brighter than the median of each measurement as the response variable and the presence of *tpn*, *iee* and the interaction of the two as the treatments. We found that the synergy of *tpn* and *iee* significantly facilitated the appearance of cells with intense fluorescence (*F*_1,12_ = 1.5 × 10^−3^, *P* = 1.0 × 10^−10^, coefficient = 3.8 × 10^−2^). Because the combined expression of *tpn* and *iee* has been associated with the excision of IS*3* (25), this suggests that, in the bright cells, IS*3* was excised, and *yfp* and *rfp* were transcribed from the stronger promoter of *kanR*.

**Figure 3.**
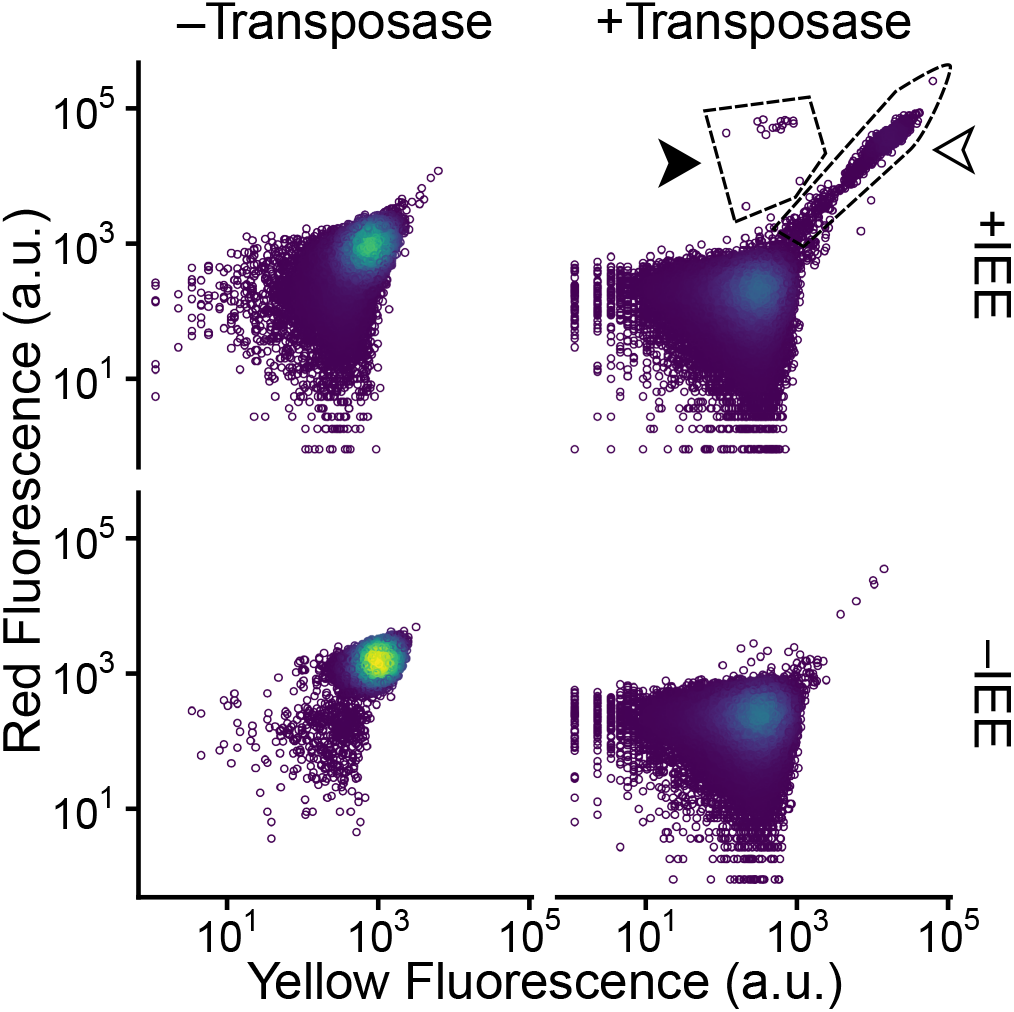
Single-cell fluorescence of cells with and without *tpn* and *iee* measured by flow cytometry (n = 50,000). The colors of the points indicate the density of points in the fraction; yellow indicates dense, and blue indicates sparse.

Besides cells showing both stronger yellow and red fluorescence, some cells with *tpn* and *iee* showed stronger red fluorescence but modest yellow fluorescence (Figure 3, black arrowhead). This implies that, as intended, not just the excision of IS*3* but also the deletion of adjacent *yfp* may have occurred.

We also noticed that the intensity of fluorescence decreased with the expression of either or both of *tpn* and *iee* (Figure 3, Supplementary Figure 2.A). ANOVA with the median log fluorescence values as the response variable and the expression of *tpn, iee*, and the interaction of the two as treatments showed that the influence of *tpn* expression was the largest and most significant (*F*_1,12_ = 7.9 × 10^2^, *P* = 2.6 × 10^−12^), corresponding to a 0.15-fold decrease in intensity. The expression of *iee* reduced the intensity by 0.64-fold, and the influence was also significant (*F*_1,12_ = 4.3 × 10, *P* = 2.6 × 10^−5^). Nevertheless, no apparent growth defect was observed (Supplementary Figure 2.B), and the fluorescent protein genes of cells collected from the major fraction of the distributions were not deleted (Supplementary Figure 3). The unexpected decrease in fluorescence seems to indicate that, during operon formation, the activity of IS can alter the expression pattern of neighboring genes.

### IS generated a variety of genotypes, including those that seem to have formed operons

To determine whether the altered fluorescence of the cells reflects the formation of novel operons, some cells with *tpn* and with or without *iee* were sorted based on their fluorescence (gates P1-3) by FACS (Figure 4.A). First, cells with both *tpn* and *iee* that evolved as four independent populations were analyzed. Consistent with Figure 3, a notable proportion of cells were found in P2 (*p* = 10^−1.3±0.1^, CI), and a few cells in P1 and P3 were also found with probabilities of 10^−4.1±0.4^ (CI) and 10^−3.3±0.4^ (CI), respectively. To investigate the diverse genotypes that seemed to have led to the formation of *kanR-rfp* operons, cells in P1 were collected from all four independent cultures. Cells in P2 and P3 were also collected from one population of cells. Next, a population of cells without the *iee* was also analyzed. Although the probability was negligible (*p* < 10^−5.5^), cells in gate P1 were also collected for further analysis (Figure 4.B).

**Figure 4.**
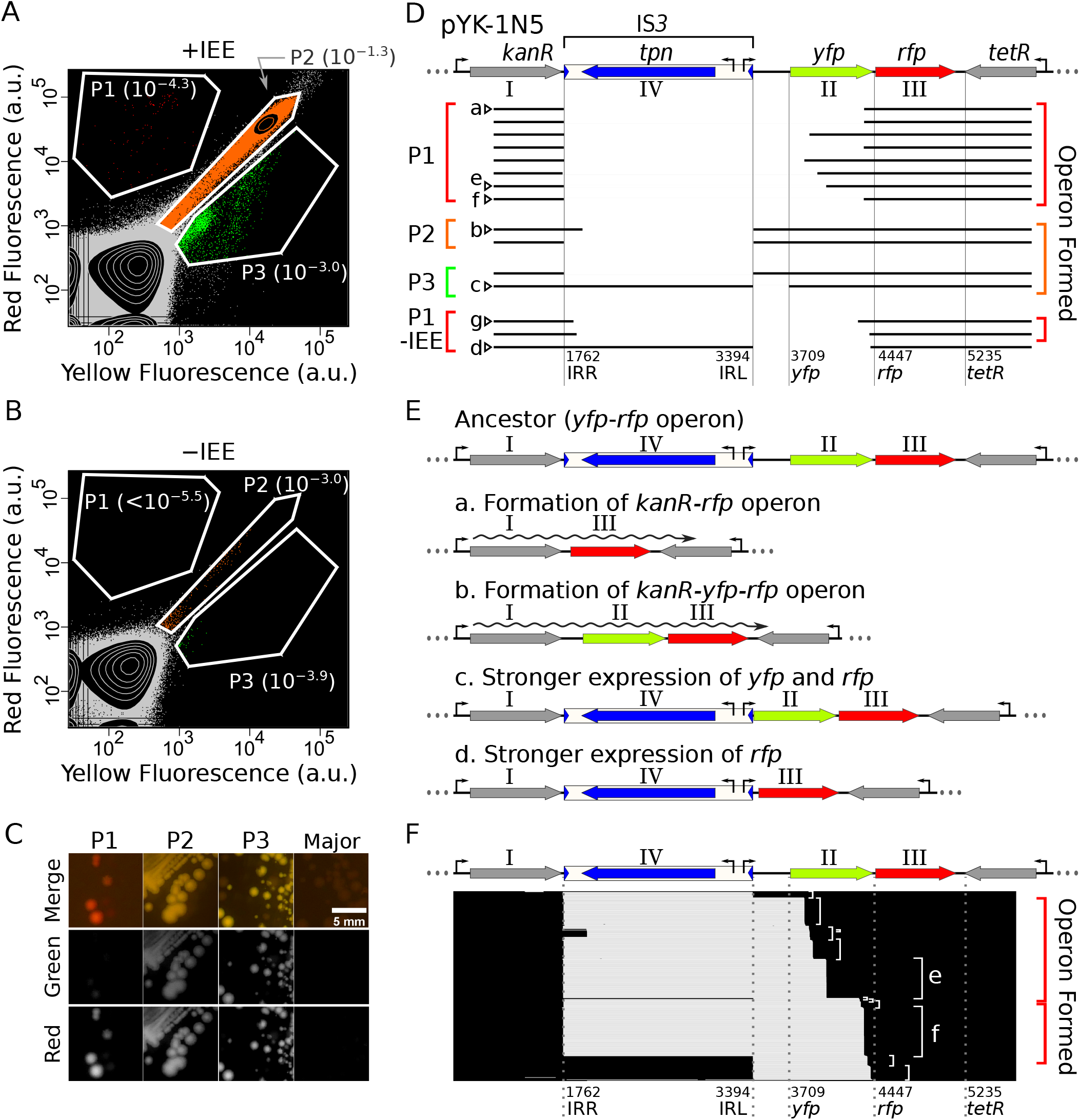
(A, B) Red and yellow fluorescence of cells with the IS*3* transposase and with (A) or without (B) insertion sequence excision enhancer (IEE) expression vectors after evolution overnight (A: n = 1,986,386; B: n = 300,644). The transcription of both transposase and IEE genes (when present) were fully induced. Each polygon (P) indicates the gates used to sort cells with fluorescence-activated cell sorting (FACS) for further analysis. The numbers within the parentheses show the proportion of cells found within each gate. No cell was found in gate P1 of subfigure B during this measurement. The cells were prepared under the same conditions as the measurements of Figure 2.C (A: +Transposase +IEE, B: +Transpsoase –IEE) but were from an independent population of cells. (C) Colonies of cells collected by FACS under blue/green LED. (D) Typical DNA sequences of pYK-1N5 that remained after evolution were determined by Sanger sequencing and are shown as black horizontal lines. The colors of the left square brackets correspond to the colors of points in subfigures A and B. The colors of the square brackets on the right correspond to the different types of operons formed; red indicates *kanR-rfp* operon, and orange indicates *kanR-yfp-rfp* operon. The vertical lines indicate the boundaries of genetic elements. The details of each deletion are provided in Supplementary Table 2. (E) The order of genes after the deletions shown in subfigure D. Each map corresponds to the sequence with the identical characters (a–d) in subfigure D. (F) Analysis of the variation of deletions using nanopore sequencing. Plasmids extracted from a population of cells sorted from P1 in subfigure A by FACS were sequenced. The loci of deletions found in each read are shown as white horizontal lines. The white square brackets on the right indicate reads showing similar deletions.

To assess if fluorescence was heritable, the cells were spread on LB agar plates with streptomycin and kanamycin and were regrown overnight. The fluorescence of colonies under blue/green LED was largely consistent with the fluorescence measured by FCM (Figure 4.C, Supplementary Figure 4). The cells from P1 showed red fluorescence as expected; cells from P2 and P3 generally showed green to orange fluorescence. However, some cells collected from P3 did not show bright fluorescence, suggesting that gate P3 contains populations overlapping with those in the major dark fraction of cells.

Next, sequences around the IS*3* of the colonies were amplified by colony PCR to genotype the sorted cells. Fragments with deletions of typical lengths were chosen and sequenced by Sanger sequencing (Figure 4.D, Supplementary Table 2). The red colonies derived from cells with *iee* found in gate P1 showed deletions of sequences from the right inverted repeat (IRR) to *yfp*. In such cases, the plasmids tended to have the IS*3* completely deleted. This result is in line with a study (25) that showed that IEE promotes the complete excision of IS*3*. The sequenced cells collected from gate P2 had the IS*3* deleted up to the IRL. While many colonies had the IS*3* completely deleted with excision starting adjacent to the IRR, some colonies had part of the IS*3* remaining as in read b of Figure 4.D. Some cells collected from P3 had a genotype similar to those in P2, and others had only the sequence between IRL and *yfp* deleted. The latter cells had the maximum possible deletion preserving the yellow fluorescence (read c, Supplementary Figure 5), implying that diverse types of deletions were generated by the activity of the IS. Overall, these results show that in line with previous studies on IS-mediated evolution (21, 25, 30), the activity of IS causes deletions of various lengths, some of which form novel operons.

Rare cells in P1 without *iee* (Figure 4.B) also had *yfp* deleted (Figure 4.D, P1–IEE) (25). Among these cells, some cells also had the copy of IS*3* excised as in read g of figure 4.D, but with parts of IS*3* remained. This shows that although rarely, IS*3* may also be able to form operons without IEE.

The deletions found by Sanger sequencing can be categorized into four types, as exemplified by reads a–d (Figure 4.E). When sequences between IS*3* and *rfp* were deleted together with IS*3*, novel *kanR-rfp* operons formed (P1, read a). This type of deletion is consistent with the operon formation model proposed in this study. The excision of only IS*3* led to the strong promoter of *kanR* to come closer to the *yfp-rfp* operon, forming *kanR-yfp-rfp* operons (P2, read b). Some cells did not form operons but had sequences adjacent to the IS*3* deleted increasing red fluorescence. Deletion of sequences between IS*3* and *yfp* led to stronger yellow fluorescence (P3, read c), and *yfp* deletion led to the loss of yellow fluorescence (P1 –IEE, read d).

To demonstrate the diversity of operon-forming deletions IS can generate in the presence of *iee*, plasmids were extracted from cells sorted by FACS in P1. The deleted loci of the plasmids were briefly analyzed by nanopore sequencing. Consistent with the results of Sanger sequencing, deletions of various lengths were observed (Figure 4.F). The major types of deletions identified were those also found by Sanger sequencing of the colonies derived from the same population of cells collected by FACS (reads e and f). Although few, some reads had the IS*3* sequence completely preserved, similar to read d. This implies that in some cells, transposase may have been excised the IS without IEE. In addition, similar to read b, some cells seemed to have IS*3* partially left undeleted (25). These results are consistent with a previous study where some cells had intact or partially excised IS*3*, even when both IS*3* and IEE were activated (25). Although sequencing by nanopore sequence is generally error-prone, the deletions found were largely consistent with those detected by Sanger sequencing, reinforcing that plasmids collected from the population of sorted cells contained a variety of deletions, many of which have led to operon formation.

### Validation of the formation of novel operons

To validate the formation of new operons, two experiments were performed.

First, the presence of mRNA transcribed from *kanR* to *rfp* was detected by RT-PCR (Figure 5.A) using RNA extracted from cells corresponding to read e of Figure 4.D. The RNA was reverse transcribed into cDNA using the forward or reverse primer, and subsequently, the DNA was amplified using the pair of primers by PCR. We detected the expected 835 bp RNA only in the direction of *kanR* (Figure 5.B, e). This shows that the evolved cells had both *kanR* and *rfp* on the same mRNA transcribed from the promoter of *kanR*, validating the formation of a *kanR-rfp* operon. Using RNA extracted from cells corresponding to read b of Figure 4.D instead, we detected 1951 bp RNA, validating the formation of a *kanR-yfp-rfp* operon (Figure 5.B, b).

**Figure 5.**
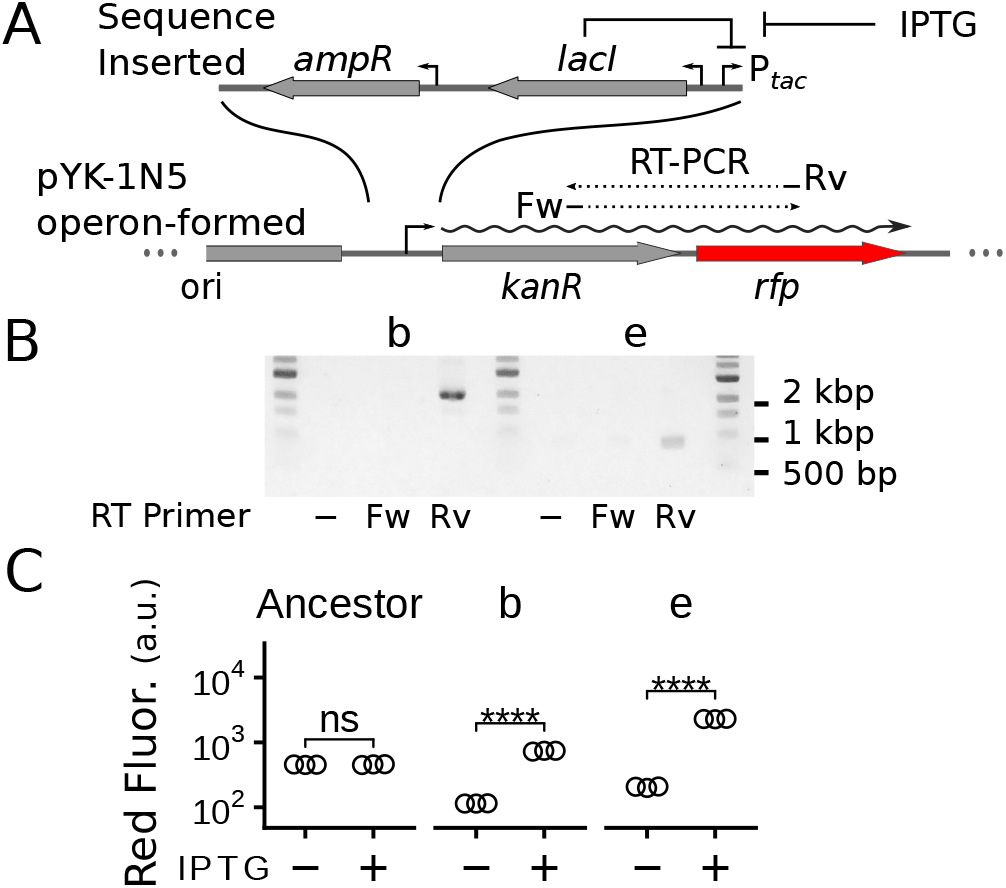
**(A)** The two experiments performed to confirm the formation of the *kanR*-*rfp* operon. **(B)** Agarose gel electrophoresis of RT-PCR products. **(C)** Median red fluorescence measured by flow cytometry of cells with re-engineered plasmids (****: *p* < 10^−4^). The constitutive promoter of *kanR* in pYK-1N5 was replaced with a DNA cassette containing an IPTG-inducible promoter (P_tac_). The lowercase letters indicate cells corresponding to reads in Figure 4.D with the same letter. Abbreviations: ampicillin resistance gene (*ampR*), lactose operon repressor gene (*lacI*), not signifiicant (ns).

Next, we tested whether the two genes could be controlled together. Specifically, we tested the coregulation of the two genes by replacing the promoter upstream of *kanR* with an IPTG inducible promoter P*_tac_* (Figure 5.A). We replaced the promoters of pYK-1N5 and its evolved derivatives and transformed the new plasmids into *E. coli* HST08 cells. The increase in the expression of *rfp* with the addition of IPTG to the growth media was checked. We found that cells with plasmid reconstructed from the ancestor plasmid did not show increased red fluorescence. In contrast, cells with plasmids reconstructed using the evolved cells showed significantly stronger fluorescence with IPTG in the growth medium (Figure 5.C). Thus, deletions by the IS allowed the coregulation of *kanR* and *rfp* as operons.

Overall, these experiments strongly support that cells evolved to have *kanR* and *rfp* to be under the control of the same promoters.

## DISCUSSION

### Summary of the results

It is widely accepted that operon structures can change dramatically (1, 9, 10, 23, 45). However, since the mechanism of operon formation has generally been studied by comparing genomes at the level above species, the very mutations that form new operons have been missed.

To explain the formation of operons, we proposed a new model whereby ISs facilitate the formation of novel operons through consecutive insertion-deletion-excision reactions (IDE model, Figure 1). According to this model, operons can form through intermediate adaptive steps (7). For experimental verification, we constructed a plasmid that mimics the first step of the proposed model (Figure 2), transformed it into an IS-less *E. coli* (Figure 3), and allowed it to evolve under the selection for kanamycin resistance and intense red fluorescence. We found that cells showed fluorescence indicative of IS-mediated deletions, especially when IEE was present (Figure 4). The structural mutations due to the transposition of IS caused rapid evolution that led to the formation of novel operons (Figure 5) overnight in 200 μL cultures.

### The IDE model is consistent with our understanding of the formation and maintenance of operons

The IDE model is consistent with how operons in prokaryotes are thought to be under selection for better gene coregulation (5, 13-15, 46) because adaptation for coregulation can begin once an operon has formed (7).

The IDE model is also consistent with our current understanding of how operons tend to form. For instance: (i) When new operons form, genes that are common across bacterial species are likely to be upstream of others (9), probably because this would preserve their sophisticated promoters even after deletion by ISs. (ii) When genes are added to existing operons, genes tend to be appended or prepended rather than inserted (9), probably because genes are added by ISs deleting redundant sequences.

### The advantages of the IDE model

The IDE model is not only consistent with previous findings but is an efficient mechanism of operon formation from three perspectives.

#### (i) Gradual formation of operons

The IDE model can incorporate the advantage of the ‘selfish operon’ model that states that operons can form without relying on rare events that directly cluster two genes into operons (7) (Figure 1.C). Even if we do not assume that higher expression of the two genes leads to higher fitness, selection for the loss of redundant sequences (12, 43, 47) coupled with ISs that delete surrounding sequences can gradually bring two genes closer, forming a new operon.

#### (ii) Formation of suboptimal operons

The expression levels of genes in operons are generally suboptimal (48), and genes in operons consisting of genes that are not functionally related tend to have functions important for cell growth (45). Our model is in line with these observations, as it explains operon formation without relying on the often-assumed selection for better gene coregulation. Consistently, various tendencies of operon formation, including the tendencies above, can be explained by assuming that operons form by selection for the preservation of genes rather than the selection for their coregulation (9, 12).

This explains an observation that inspired our new model, that is, several insect endosymbionts that have experienced extreme population bottlenecks (49) have small but operon-rich genomes (50, 51). The selection for better coregulation would be too weak in these organisms for operons to form. Because insect symbionts generally reduce their genome size through a burst and a subsequent loss of ISs (18, 52), in line with our study, ISs may have formed their operon-rich genomes.

Our model improves the ‘persistence model’ (12), which currently relies upon the selection for coregulation to explain the formation of operons (4, 13). Unlike the ‘persistence model,’ we have attributed the large tandem deletions that cluster genes to the activity of ISs (17, 18). The activity of ISs alters the expression of surrounding genes (41). This enables cells with new IS insertions to become adaptive. In addition, this ensured that the premature loss of ISs after the deletion was maladaptive (Figure 1.C). It seems that the structure of ISs that have evolved for their proliferation may have made them a source of mutation particularly suitable for the formation of operons.

#### (iii) Accelerated recombination rate

ISs not only actively transpose but are both drivers (53) and major targets of homologous recombination (54). They accelerate the formation of operons by transiently increasing the local structural mutation rates surrounding the IS (17, 30). How the increased mutation rate is kept local and transient is critical for our model because the rate of recombination can exceed the upper limit of global mutation rates set by the population size (55). In contrast, previous models based on random recombination (‘recombination model’ (11)) have been refuted because the rate of random recombination in prokaryotes seems to be too low to account for operon formation (7). Moreover, an experimental demonstration (Figure 4), unprecedented in any other models of operon formation, was possible because of the rapid mutation by IS*3*.

The accelerated rate of recombination is crucial for the IDE model to act efficiently by overcoming the assumption that all genes present between two genes are dispensable (Figure 1.C). Recombination among multiple copies of IS enables multiple pairs of beneficial genes to be tested for undergoing the deletion step of operon formation. Furthermore, within prokaryotic genomes, some loci are easier to have only dispensable genes between two beneficial genes such as: pathogenicity islands (13), plasmids (‘Scribbling Pad’ hypothesis (56)), and clusters of nonessential genes (12, 16). ISs can actively form such loci by promoting recombination. For instance, duplication by IS-mediated recombination can create redundant copies of essential genes, making them nonessential (‘SNAP hypothesis’ (19)). In addition, recombination of two copies of ISs can form plasmids (24). Plasmid pYK-1N5 (Figure 2.A) can be formed from sequences within a chromosome by homologous recombination of a pair of IS*3*.

In conclusion, the operon-rich genomes of prokaryotes may have been shaped through the coexistence of ISs in the prokaryotic genome whereby operons rapidly form by the IDE model, newly-formed operons that are beneficial acquire sophisticated regulation and get fixed in genomes (‘regulatory model’ (5)), and organisms that have acquired operon-rich genomes become robust against the activity of ISs that cause large deletions (‘persistence model’ (12, 16)).

### The activity of transposase may decrease the expression of genes surrounding an IS

An important characteristic of ISs is that they alter the expression of surrounding genes through their internal promoters (41). We observed that the increased activity of the IS*3* transposase significantly decreased the expression of neighboring genes encoding fluorescent proteins (Figure 3). However, this did not seem to be due to the deletion of *rfp* and *yfp* (Supplementary Figure 4). Moreover, no growth defect was observed (Supplementary Figure 2.B), implying that the reduced copy number of plasmids due to their excision was not the cause because pYK-1N5 is a low copy number plasmid, and decreasing its copy number should have been detrimental to growth.

Rather, we speculate that the decreased fluorescence could have been due to the nicking of DNA by the transposase, the binding of transposase to the IRs, the formation of a protein DNA complex called ‘Figure-eight’ (40), or the increased transcription itself strengthening the supercoiling of DNA (57). While solving the dominant cause of the decreased fluorescence is out of our scope, we speculate that ISs may act as global regulators, with IS transposase acting as transcription repressors and IS inverted repeats as operator sequences. Supporting this, ISs have been found in orientations that interfere with neighboring genes in clinical samples of *Staphylococcus aureus* (58).

### Limitations of our study

Our model is limited in that some operons are likely to have formed using mechanisms other than that proposed by our model. For instance, new operons can form by replacing the genes of preexisting operons (9) or by duplication (10). Also, toxin-antitoxin systems form operons, probably by the ‘selfish operon’ model.

Second, although our experiment shows that ISs cause diverse deletions that can form operons, the process of operon formation may likely be much slower in nature without the enhanced activity of the IS and IEE. Consistently, in our experimental demonstration, the rate of operon formation was much lower when IEE was not present (Figure 4.B, P1). Nevertheless, as our observations were largely consistent with those of previous studies, we believe that the results are robust to minor changes in the design of the experimental model.

Finally, while deletion and excision of IS is common in major types of ISs (20, 22), we only demonstrated the formation of operons using the combination of IS*3* and IEE in a strain of *E. coli*. We believe, to extend the scope of our model, it is necessary to understand the genetic factors essential for the insertion, deletion, and excision reactions by IS transposase and the types of ISs that promote operon formation. Future studies in such directions might also reveal the origins of ancient operons (10, 19) by comparing when the genetic factors, the ISs, and the operons emerged.

### Future work

Our study is probably the first experimental demonstration of operon formation in the laboratory. However, the proposed model requires validation in nature. This is important as ISs are regarded as drivers of operon destruction (19, 23), although this view may be biased because studies have focused more on the evolution of conserved operons than on their formation.

A potential difficulty to find examples of operon formation in nature is that IS can be thought of as a genetic “catalyst” that locally accelerates the mutation rate and transforms a genome into a new genome with an additional operon (Figure 1.C). As a catalyst, the activity of IS is virtually traceless. As a result, previous studies have overlooked the role of deletions by IS in operon formation because, based on parsimony of events, operon formation by the IDE model cannot be distinguished from large tandem deletions without ISs (9). Indeed, partial sequences of ISs may remain as read g of Figure 4.D, but these sequences would be rapidly lost, as they are unlikely to have any function. We believe that to find cases where operons are formed by our model and to determine whether ISs are the major drivers of operon formation, analyzing ongoing evolutions in detail (28, 54, 59, 60) to detect the intermediate steps of operon formation is essential.

We postulate that these steps may potentially be identified in organisms that are under reductive evolution (61), which ISs are known to facilitate (18, 43). When genomes expand, ISs can potentiate newly acquired genes to form operons by activating them. As genome size decreases, ISs delete redundant sequences and excise themselves, forming new operons. Supporting this idea, a systematic study of genome size evolution in cyanobacteria showed that operon-rich genomes tend to have experienced genome reduction (14). In contrast, larger genomes with many ISs are relatively poor in operons (4), perhaps because they are yet to experience an upsurge in operon formation. Future studies on, for example, the evolution of pathogenic *E. coli* with multiple ISs that degrade their genomes (62) and are excised by IEE (63), or organisms with genome size evolution artificially accelerated using hyper-active transposons (64) may provide a clearer picture of the coevolution of prokaryotic genomic and regulatory architecture with ISs, as suggested by our study.

## DATA AVAILABILITY

Nanopore sequencing data have been deposited with SRA (BioProject PRJNA768397).

The raw flow cytometry data are available in FlowRepository (FR-FCM-Z4LV).

## SUPPLEMENTARY DATA

Supplementary Data are available at NAR Online: Supplementary Figures 1–6, Supplementary Tables 1–2, and DNA sequences.

## AUTHOR CONTRIBUTIONS

Conceptualization and methodology: Y.K. and S.T.; data curation, formal analysis, investigation, validation, and visualization: Y.K.; funding acquisition: Y.K., S.T., and C.F.; project administration: S.T.; resources and supervision: S.T. and C.F.; writing – original draft: Y.K.; writing – review & editing: Y.K., S.T., and C.F.

## ACKNOWLEDGMENTS

The authors thank the following people for generously sharing their reagents: The mScarlet-I sequence was generated by mutating the mScarlet sequence, which was a gift from Dr. Ryudo Ohbayashi. The Tn*5* transposase sequence used as –*tpn* control was a gift from Dr. Ronald Chalmers. The Marionette Sensor Collection was a gift from Dr. Christopher Voigt (Addgene #1000000137). pCDSSara-L18 was a gift from Dr. Peter Schultz.

## FUNDING

This work was supported by the Japan Society for the Promotion of Science [21J20693 to Y.K., 18H02427 to S.T., 17H06389 to C.F, 19H05626 to C.F]; and the Japan Science and Technology Agency [JPMJER1902 to C.F]. Funding for open access charge: the Japan Science and Technology Agency.

## Conflict of interest statement

None declared.

## Notes

### Competing Interest Statement

The authors have declared no competing interest.

