## Supplementary material for "Operon formation by insertion sequence IS*3* in *Escherichia coli*": Online Supplementary Information

**Supplementary Information of "Operon formation by insertion sequence IS*3* in *Escherichia coli*"**

### Supplementary Figures

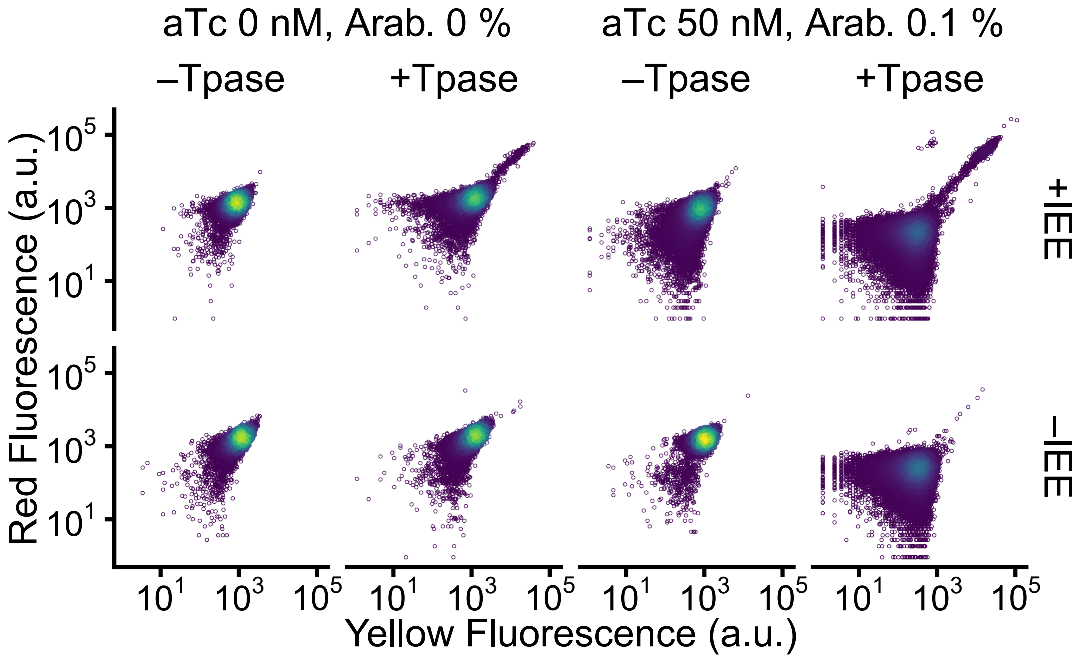


**Supplementary Figure 1:** Single-cell fluorescence of cells with and without *tpn* and *iee* measured by flow cytometry. After subsetting the measurements with forward scatter values within a two-fold range, 50,000 measurements were randomly sampled for every condition. The colors of the points indicate the population density in the fraction, with yellow being dense and blue being sparse.

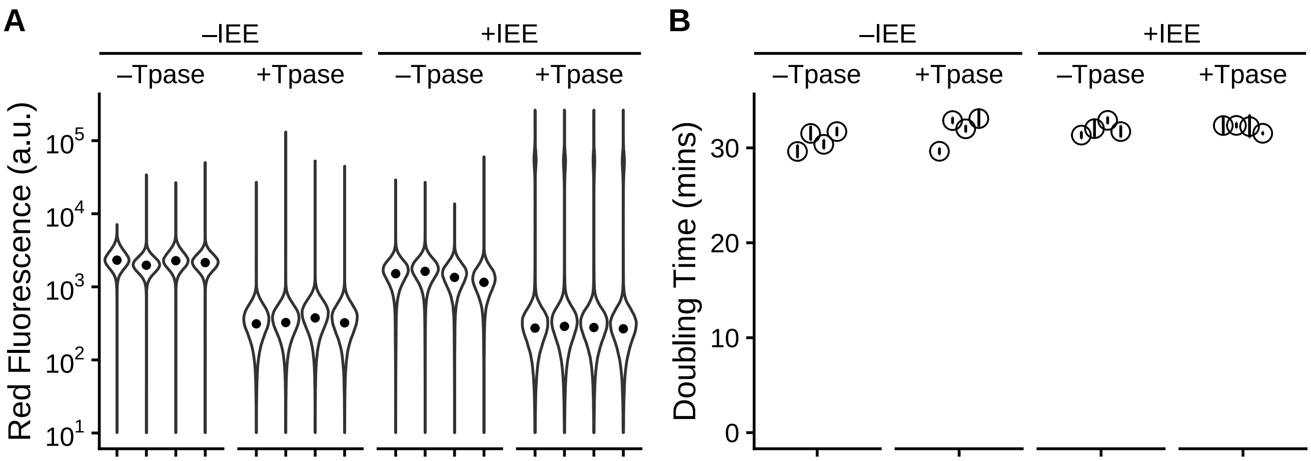


**Supplementary Figure 2: (A)** Red fluorescence measured by FCM. The median red fluorescence measurements used for ANOVA are indicated as black dots. **(B)** The doubling time of cells. The error bars indicate the 95 % confidence intervals assuming normality of errors in the relative log turbidity.

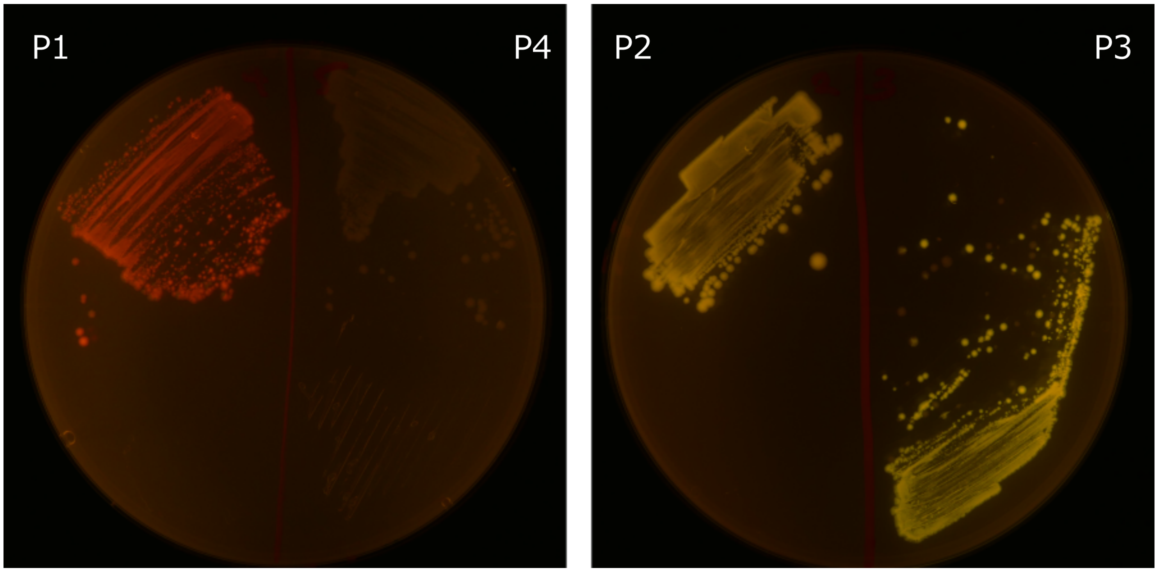


**Supplementary Figure 3:** Fluorescence of colonies of cells sorted by FACS. Cells sorted by FACS were cultured overnight in liquid media and streaked onto agar plates for colony PCR. P4 indicates the gate used to sort the center of the major fraction of cells.

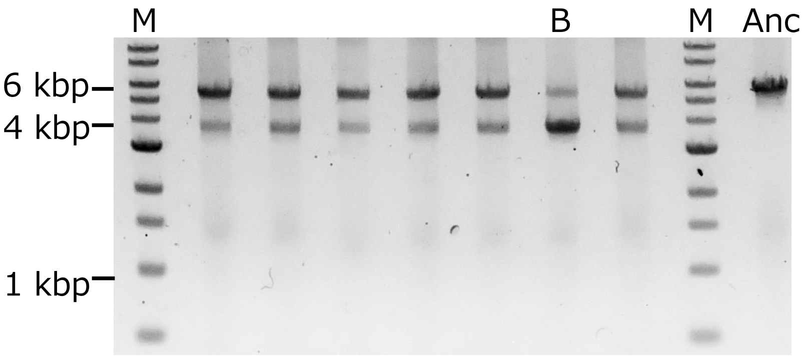


**Supplementary Figure 4:** Colony PCR of cells in the major fraction of dark +*iee* +*tpn* cells collected by FACS. Sanger sequencing of the longer bands was performed, confirming that the full 5423 bp of the sequence between ori and *kanR* was preserved as was in the ancestor. Some of the collected colonies had bright fluorescence. This colony is marked as *B*. How all the colonies had a second short band identical to *B* indicates that, within the colonies, some cells had the IS*3* excised. M: marker (1 kbp ladder), B: bright cells, Anc: ancestor.

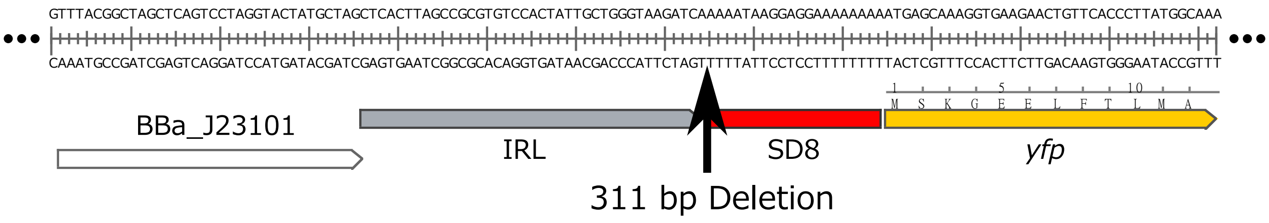


**Supplementary Figure 5:** Deletion of sequences up to the ribosomal binding site (SD8) upstream *yfp*.

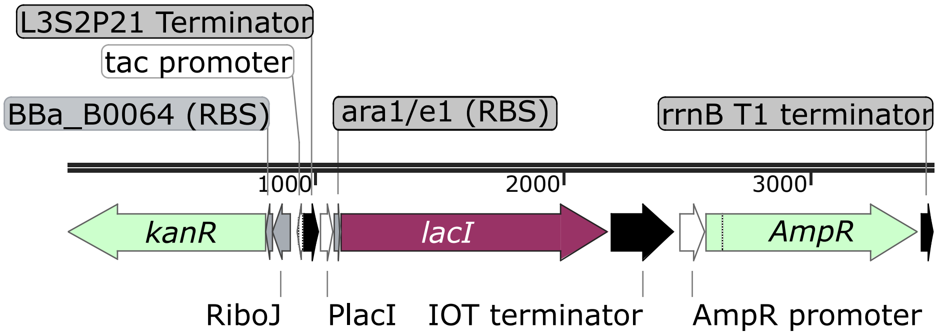


**Supplementary Figure 6:** Detailed map of sequences placed upstream of *kanR* to make *kanR* inducible with IPTG. The sequence including this region is provided in the *DNA Sequences* section.

### Supplementary Tables

**Supplementary Table 1:** Primers used in this study.

| | Forward | Reverse |
|---|---|---|
| **Figure 4** | | |
| P1 (Sanger seq.) | GCTTTACGGTATCGCCGCTC | GTTCATGGAACCTTCCATGTG |
| P2, 3 (Sanger seq.) | GCTTTACGGTATCGCCGCTC | CATTCACGTCACCATCCAGC |
| **Figure 5** | | |
| RT-PCR | GGCGCCACCGTCTTCAAAAT | GACCACCAAGCGAAACATCG |
| In-Fusion Vector | GTTGTTTGTCGGTGAACTGTAAGACGGGTAAGCCTGTTGATGA | AGGGGAAATACTAGATGATGATTGAACAAGATGGATTGCACGCA |
| In-Fusion Insert | CATCTAGTATTTCCCCTCTTTCTCTAGTATTAAACAA | CAGTTCACCGACAAACAACAGATAA |
| **Supplementary Figure 4** | | |
| Major Dark Fraction | GCTTTACGGTATCGCCGCTC | TGTGTGGCACTACTCAACCC |

**Supplementary Table 2:** The sequences of pYK-1N5 that were deleted. The genotypes of cells collected by FACS under the gates P1-3 were determined by sanger sequencing. The reads numbered in column *Figure 4.D* correspond to the representative reads shown in the figure. The solid and pale letters in the *Deletion* column indicate the ends of preserved and deleted sequences, respectively. The locus numbers of the deleted sequences correspond to the numbers found in the sequence of PYK-1N5 provided in the *DNA Sequences* section.

| Gate | Population | Figure 4.D | From | To | Deletion (bps) |
|---|---|---|---|---|---|
| P1 | 1 | 1 read a | 1761 | 4358 | TGGGG TTCTG 2598 AAGAC CCGAA |
| P1 | 1 | 2 | 1761 | 4358 | TGGGG TTCTG 2598 AAGAC CCGAA |
| P1 | 1 | | 1761 | 4358 | TGGGG TTCTG 2598 AAGAC CCGAA |
| P1 | 1 | | 1761 | 4358 | TGGGG TTCTG 2598 AAGAC CCGAA |
| P1 | 1 | | 1761 | 4358 | TGGGG TTCTG 2598 AAGAC CCGAA |
| P1 | 2 | 3 | 1764 | 3885 | GGTTC TGATC 2122 CAAAC TGCCT |
| P1 | 2 | | 1764 | 3885 | GGTTC TGATC 2122 CAAAC TGCCT |
| P1 | 2 | 4 | 1761 | 4355 | TGGGG TTCTG 2595 CTAAA GACCC |
| P1 | 3 | 5 | 1764 | 3841 | GGTTC TGATC 2078 GCTAC TTATG |
| P1 | 3 | 6 | 1751 | 3954 | AGCGG GACTC 2204 CCCGG ATCAC |
| P1 | 4 | 7 read e | 1761 | 4030 | TGGGG TTCTG 2270 TTCAA GGACG |
| P1 | 4 | 8 read f | 1761 | 4355 | TGGGG TTCTG 2595 CTAAA GACCC |
| P2 | 1 | 1 read b | 1923 | 3401 | GAGGT TCTTG 1479 AGCTG GCACT |
| P2 | 1 | 2 | 1761 | 3402 | TGGGG TTCTG 1642 GCTGG CACTG |
| P2 | 1 | | 1767 | 3402 | TCTGA TCCTA 1636 GCTGG CACTG |
| P3 | 1 | 1 | 1761 | 3400 | TGGGG TTCTG 1640 CAGCT GGCAC |
| P3 | 1 | | 1761 | 3400 | TGGGG TTCTG 1640 CAGCT GGCAC |
| P3 | 1 | 2 read c | 3398 | 3708 | GATCA GCTGG 311 TGGAG AAAAT |
| P3 | 1 | | 3398 | 3708 | GATCA GCTGG 311 TGGAG AAAAT |
| P3 | 1 | | 3398 | 3708 | GATCA GCTGG 311 TGGAG AAAAT |
| P1 | −IEE | 1 read g | 1845 | 4305 | TCGGG AGGCC 2461 CCCAG TACTG |
| P1 | −IEE | 2 | 1872 | 4404 | TACCG AGTCC 2533 TACCG CGGCA |
| P1 | −IEE | 3 read d | 3398 | 4413 | GATCA GCTGG 1016 AGGCA TTACC |

### DNA Sequences

pYK-1N5 (Fluorescence reporter vector)

```
   1 actactttag tcagttccgc agtattacaa aaggatgtcg caaacgctgt ttgctcctct
  61 acaaaacaga ccttaaaacc ctaaaggctt aagtagcacc ctcgcaagct cgggcaaatc
 121 gctgaatatt ccttttgtct ccgaccatca ggcacctgag tcgctgtctt tttcgtgaca
 181 ttcagttcgc tgcgctcacg gctctggcag tgaatggggg taaatggcac tacaggcgcc
 241 ttttatggat tcatgcaagg aaactaccca taatacaaga aaagcccgtc acgggcttct
 301 cagggcgttt tatggcgggt ctgctatgtg gtgctatctg actttttgct gttcagcagt
 361 tcctgccctc tgattttcca gtctgaccac ttcggattat cccgtgacag gtcattcaga
 421 ctggctaatg cacccagtaa ggcagcggta tcatcaacag gcttacccgt cttactgtca
 481 attcttgaag acgaaagggc ctcgtgatac gcctattttt ataggttaat gtcatgataa
 541 taatggtttc ttagacgtca ggtggcactt ttcggggaaa tgtgcgcgga acccctattt
 601 gtttattttt ctaaatacat tcaaatatgt atccgctcat gagacaataa ccctgataaa
 661 tgcttcaata atattgaaaa aggaagagta tgagtattca acatttccgt gtcgccctta
 721 ttcccttttt tgcggcattt tgccttcctg tttttgctca cccagaaacg ctggtgaaag
 781 taaaagatgc tgaagatcag ttgggtgcac gagtgggtta catcgaactg gatctcaaca
 841 gcggtaagat ccttgagagt tttcgccccg aagaacgctc atgtttgaca gcttatcatc
 901 gatatgcttt aatgcggtag tgatcaagag acaggatgag gatcgtttcg catgattgaa
 961 caagatggat tgcacgcagg ttctccggcc gcttgggtgg agaggctatt cggctatgac
1021 tgggcacaac agacaatcgg ctgctctgat gccgccgtgt tccggctgtc agcgcagggg
1081 cgcccggttc tttttgtcaa gaccgacctg tccggtgccc tgaatgaact gcaggacgag
1141 gcagcgcggc tatcgtggct ggccacgacg ggcgttcctt gcgcagctgt gctcgacgtt
1201 gtcactgaag cgggaaggga ctggctgcta ttgggcgaag tgccggggca ggatctcctg
1261 tcatctcacc ttgctcctgc cgagaaagta tccatcatgg ctgatgcaat gcggcggctg
1321 catacgcttg atccggctac ctgcccattc gaccaccaaa cgaaacatcg catcgagcga
1381 gcacgtactc ggatggaagc cggtcttgtc gatcaggatg atctggacga agagcatcag
1441 gggctcgcgc cagccgaact gttcgccagg ctcaaggcgc gcatgcccga cggcgaggat
1501 ctcgtcgtga cccatggcga tgcctgcttg ccgaatatca tggtggaaaa tggccgcttt
1561 tctggattca tcgactgtgg ccggctgggt gtggcggacc gctatcagga catagcgttg
1621 gctacccgtg atattgctga agagcttggc ggcgaatggg ctgaccgctt cctcgtgctt
1681 tacggtatcg ccgctcccga ttcgcagcgc atcgccttct atcgccttct tgacgagttc
1741 ttctgagcgg gactctgggg ttctgatcct acccacgtaa tatggacaca ggcctaagcg
1801 agcgccgcca ttggaccaaa acgaaaaaag gcccccccttt cgggaggcct cttttctgga
1861 atttggtacc gagtccatac acgtttttttg ggctaactat ttgttaactg tctaagcgag
1921 gttcttgttt tcaaattgtt ccggactgag gccgccacac caactgtgcc gccgccaccg
1981 attgtaatca cattcgatat aattaaacac cgttgcccgc attatttccc ggctgataaa
2041 gtgttctcca tggatacatt ccactttcag cgaatgaaag aagctttcca cgcaggcatt
2101 atcgtagcag caacctttttg cgctcatact tccacgcaga ttatgccgct tcagttgcgc
2161 ctgataatct gctgaacagt actggcctcc acggtccgtg tgaacgataa cgttccgggg
2221 cctcttacgc cgccacagcg ccatctgcag ggcatcgcag gccagttgcg ccgtcatgcg
2281 tggcgacatt gaccagccaa taacggcacg tgaccacagg tcaatgacca ctgccagata
2341 cagccagcct tcatctgtac gtaagtacgt gatgtctcct gcccacttct ggttcgggcc
2401 actggcgtaa aaatcctgct ccaacagatt ttctgacaca ggcaggccgt gtgcgcggta
2461 gctgaccggg ctgaacttcc gggaggcctt tgccctcagt ccctgacggc gcaggcttgc
2521 cgccacggtt tttacgttaa aggggtaacc ctgagcacgc agttcatccg tcaggcgtgg
2581 ggcaccgtaa cgctgttttg accgggtaaa agccgcgagg acaacgctgt cgcagtgttg
2641 gcggaactgc tgacgcgtgc ttatccttgt ccgccgctga caccacgtat accagccgct
2701 gcgagccacc cggagcacgc ggcacattgc tttgatgctg aactcagcct gatgtttttc
2761 aataaagaca tacttcattt caggcgcttc gcgaagtatg tcgcggccct tttggaggat
2821 agccagctct tcatcccgtt ctgccagctg gcgtttgaga cgtgcaatct cggtagacat
2881 ctccagttca cgttcagaag acgtctgctg attttgctgt ttactgcgcc agttgtagag
2941 ttgtgattca tacaggctga gttcacgggc tgcggcagta acaccgatgc gttcagcaag
3001 cttcagggct tcactgcgaa attcaggcga atgctgttta cggggttttt tactggttga
3061 tactgttttt gtcatctagt atttcccctc tttctctagt attaaacaaa attatttgta
3121 gaggctgttt cgtcctcacg gactcatcag accggaaagc acatccggtg acagctgtgc
3181 tcattatatc tatcactgat agggatgtca atctctatca ctgataggga ggtcagtgcg
3241 tcctgctgaa aacagccaac gatcgttaag cgggagacca gaaacaaaaa aaggcccccc
3301 gttagggagg ccttcaataa ttggtttacg gctagctcag tcctaggtac tatgctagct
```

3361 cacttagccg cgtgtccact attgctgggt aagatcagct ggcacttgcg tataggtcac
3421 ctcggcagaa atagcatgcg ggtctttctg gcgcttatct ttaccaaaca gccccgacttc
3481 atcgccataa tactgttcat acatcaggct tgcgccaagc tgcggccagg cgggtaaata
3541 gccctcagca cgaatatccc agccattcgc cgggcgttcc tgataatcct caatatccgg
3601 cgattttttc cagccagaag cccgaatata accattggcg ctcagtttca gataatcgcg
3661 ccagtattcc gcaccaacac caatgcgggt atgaagatct tcctggagaa aataaggagg
3721 aaaaaaaaat gagcaaaggt gaagaactgt tcaccggcgt tgtgccaatt ctggttgagc
3781 tggatggtga cgtgaatggc cacaaatttt ccgtgtctgg tgaaggcgag ggtgatgcta
3841 cttatggcaa actgactctg aaactgatct gtaccaccgg caaactgcct gttccgtggc
3901 caactctggt cactactctg ggttacggcc tgatgtgttt tgcgcgttac ccggatcaca
3961 tgaaacagca tgacttcttc aaatctgcca tgccggaagg ctatgtccaa gaacgtacga
4021 tctttttcaa ggacgacggc aactataaaa cccgtgccga agttaaattc gagggtgaca
4081 ccctggttaa ccgcatcgaa ctgaaaggca ttgacttcaa agaggacggc aacattctgg
4141 gtcacaagct ggaatacaac tacaactccc acaacgttta cattactgct gacaagcaga
4201 aaaacggcat caaagcaaac ttcaagatcc gtcacaacat tgaagatggt ggcgtacagc
4261 tggcagatca ctaccagcag aacactccaa tcggtgatgg cccagtactg ctgccagata
4321 accattacct gtcctaccag agcaaactgt ctaaagaccc gaacgaaaaa cgtgaccaca
4381 tggtactgct ggaatttgtt accgcggcag gcattaccca cggtatggac gaactgtata
4441 aataataaaa taaggaggaa aaaaaatggt gtccaagggc gaggctgtca ttaaggagtt
4501 catgcgcttt aaagtccaca tggaaggttc catgaacggc cacgagttcg aaattgaggg
4561 cgagggtgaa ggtcgcccct acgaaggcac tcagaccgcc aaattaaagg tgaccaaagg
4621 cgggccgctc ccctttagtt gggatatcct cagtccccag ttcatgtacg ggagtcgggc
4681 cttcattaag cacccggctg atattcccga ctactataaa caatcctttc cggaaggttt
4741 taagtgggaa cgtgtgatga attttgaaga cggtggcgcc gtaaccgtca cacaagacac
4801 aagcctcgaa gacggcaccc tgatttataa ggtaaaactg cgtggcacca actttccacc
4861 tgacggcccc gtcatgcaaa aaaaaacgat gggctgggaa gcctctaccg aacggctgta
4921 tcctgaagat ggagtgttga aaggtgatat caagatggca ctccggttaa aagatggggg
4981 gcgctacttg gcggatttta aaaccactta caaagcgaaa aaacccgtgc agatgccagg
5041 agcgtataat gttgatcgca aattggatat cacgtcgcat aatgaagatt atacggtggt
5101 tgaacaatat gaacgctctg aaggacggca ttcgaccggg gggatggatg aactttataa
5161 ataagtcaca ctggctcacc ttcgggtggg cctttctgcg tttaattaac agttaacaat
5221 cagtaggatc cttatcagga cccactttca catttaagtt gtttttctaa tccgcatatg
5281 atcaattcaa ggccgaataa gaaggctggc tctgcacctt ggtgatcaaa taattcgata
5341 gcttgtcgta ataatggcgg catactatca gtagtaggtg tttccctttc ttctttagcg
5401 acttgatgct cttgatcttc caatacgcaa cctaaagtaa aatgccccac agcgctgagt
5461 gcatataatg cattctctag tgaaaaacct tgttggcata aaaaggctaa ttgattttcg
5521 agagtttcat actgtttttc tgtaggccgt gtacctaaat gtacttttgc tccatcgcga
5581 tgacttagta aagcacatct aaaactttta gcgttattac gtaaaaaatc ttgccagctt
5641 tccccttcta aagggcaaaa gtgagtatgg tgcctatcta acatctcaat ggctaaggcg
5701 tcgagcaaag cccgcttatt ttttacatgc caatacaatg taggctgctc tacacctagc
5761 ttctgggcga gtttacgggt tgttaaacct tcgattccga cctcattaag cagctctaat
5821 gcgctgttaa tcactttact tttatctaat ctggacatat tcaccaccct gaattgactc
5881 tcttccgggc gctatcatgc cataccgcga aaggttttgc gccattcgat ggcgcgccgc
5941 tgtaacaagt tgtctcaggt gttcaatttc atgttctagt tgctttgttt tactggtttc
6001 acctgttcta ttaggtgtta catgctgttc atctgttaca ttgtcgatct gttcatggtg
6061 aacagctttg aatgcaccaa aaactcgtaa aagctctgat gtatctatct tttttacacc
6121 gttttcatct gtgcatatgg acagttttcc ctttgatatg taacggtgaa cagttgttct
6181 acttttgttt gttagtcttg atgcttcact gatagataca agagccataa gaacctcaga
6241 tccttccgta tttagccagt atgttctcta gtgtggttcg ttgtttttgc gtgagccatg
6301 agaacgaacc attgagatca tacttacttt gcatgtcact caaaaatttt gcctcaaaac
6361 tggtgagctg aatttttgca gttaaagcat cgtgtagtgt tttttcttagt ccgttatgta
6421 ggtaggaatc tgatgtaatg gttgttggta ttttgtcacc attcattttt atctggttgt
6481 tctcaagttc ggttacgaga tccatttgtc tatctagttc aacttggaaa atcaacgtat
6541 cagtcgggcg gcctcgctta tcaaccacca atttcatatt gctgtaagtg tttaaatctt
6601 tacttattgg tttcaaaacc cattggttaa gccttttaaa ctcatggtag ttattttcaa
6661 gcattaacat gaacttaaat tcatcaaggc taatctctat atttgccttg tgagttttct
6721 tttgtgttag ttcttttaat aaccactcat aaatcctcat agagtatttg ttttcaaaag
6781 acttaacatg ttccagatta tattttatga attttttttaa ctggaaaaga taaggcaata
6841 tctcttcact aaaaactaat tctaattttt cgcttgagaa cttggcatag tttgtccact

```
6901 ggaaaatctc aaagccttta accaaaggat tcctgatttc cacagttctc gtcatcagct
6961 ctctggttgc tttagctaat acaccataag cattttccct actgatgttc atcatctgag
7021 cgtattggtt ataagtgaac gataccatcc gttctttcct tgtagggttt tcaatcgtgg
7081 ggttgagtag tgccacacag cataaaatta gcttggtttc atgctccgtt aagtcatagc
7141 gactaatcgc tagttcattt gctttgaaaa caactaattc agacatacat ctcaattggt
7201 ctaggtgatt ttaatcacta taccaattga gatgggctag tcaatgataa ttactagtcc
7261 tttttctttg agttgtgggt atctgtaaat tctgctagac ctttgctgga aaacttgtaa
7321 attctgctag accctctgta aattccgcta gacctttgtg tgtttttttt gtttatattc
7381 aagtggttat aatttataga ataaagaaag aataaaaaaa gataaaaaga atagatccca
7441 gccctgtgta taactc
```
//

pYK-1S3 (IEE expression vector)

```
   1 ttatttgtta actgttaatt gtccttgttc aaggatgctg tctttgacaa cagatgtttt
  61 cttgcctttg atgttcagca ggaagctcgg cgcaaacgtt gattgtttgt ctgcgtagaa
 121 tcctctgttt gtcatatagc ttgtaatcac gacattgttt cctttcgctt gaggtacagc
 181 gaagtgtgag taagtaaagg ttacatcgtt aggatcaaga tccattttta acacaaggcc
 241 agttttgttc agcggcttgt atgggccagt taaagaatta gaaacataac caagcatgta
 301 aatatcgtta gacgtaatgc cgtcaatcgt catttttgat ccgcgggagt cagtgaacag
 361 gtaccatttg ccgttcattt taaagacgtt cgcgcgttca atttcatctg ttactgtgtt
 421 agatgcaatc agcggtttca tcactttttt cagtgtgtaa tcatcgttta gctcaatcat
 481 accgagagcg ccgtttgcta actcagccgt gcgtttttta tcgctttgca gaagtttttg
 541 actttcttga cggaagaatg atgtgctttt gccatagtat gctttgttaa ataaagattc
 601 ttcgccttgg tagccatctt cagttccagt gtttgcttca aatactaagt atttgtggcc
 661 tttatcttct acgtagtgag gatctctcag cgtatggttg tcgcctgagc tgtagttgcc
 721 ttcatcgatg aactgctgta cattttgata cgtttttccg tcaccgtcaa agattgattt
 781 ataatcctct acaccgttga tgttcaaaga gctgtctgat gctgatacgt taacttgtgc
 841 agttgtcagt gtttgtttgc cgtaatgttt accggagaaa tcagtgtaga ataaacggat
 901 ttttccgtca gatgtaaatg tggctgaacc tgaccattct tgtgtttggt cttttaggat
 961 agaatcattt gcatcgaatt tgtcgctgtc tttaaagacg cggccagcgt tttccagct
1021 gtcaatagaa gtttcgccga ctttttgata gaacatgtaa atcgatgtgt catccgcatt
1081 tttaggatct ccggctaatg caaagacgat gtggtagccg tgatagtttg cgacagtgcc
1141 gtcagcgttt tgtaatggcc agctgtccca aacgtccagg ccttttgcag aagagatatt
1201 tttaattgtg gacgaatcaa attcaggaac ttgatatttt tcattttttt gctgttcagg
1261 gatttgcagc atatcatggc gtgtaatatg ggaaatgccg tatgtttcct tatatggctt
1321 ttggttcgtt tctttcgcaa acgcttgagt tgcgcctcct gccagcagtg cggtagtaaa
1381 ggttaatact gttgcttgtt ttgcaaactt tttgatgttc atcgttcatg tctccttttt
1441 tatgtactgt gttagcggtc tgcttcttcc agccctcctg tttgaagatg gcaagttagt
1501 tacgcacaat aaaaaaagac ctaaaatatg taaggggtga cgccaaagta tacactttgc
1561 cctttacaca ttttaggtct tgcctgcttt atcagtaaca aacccgcgcg atttactttt
1621 cgaactcatt ctattagact ctcgtttgga ttgcaactgg tctattttcc tcttttgttt
1681 gatagaaaat cataaaagga tttgcagact acgggcctaa agaactaaaa aatctatctg
1741 tttcttttca ttctctgtat tttttatagt ttctgttgca tgggcataaa gttgcctttt
1801 taatccctag gaataggaac ttcggaatga gctcttattt gccgactacc ttggtgatct
1861 cgcctttcac gtagtggaca aattcttcca actgatctgc gcgcgaggcc aagcgatctt
1921 cttcttgtcc aagataagcc tgtctagctt caagtatgac gggctgatac tggccggca
1981 ggcgctccat tgcccagtcg gcagcgacat ccttcggcgc gattttgccg gttactgcgc
2041 tgtaccaaat gcgggacaac gtaagcacta catttcgctc atcgccagcc cagtcgggcg
2101 gcgagttcca tagcgttaag gtttcattta gcgcctcaaa tagatcctgt tcaggaaccg
2161 gatcaaagag ttcctccgcc gctggaccta ccaaggcaac gctatgttct cttgcttttg
2221 tcagcaagat agccagatca atgtcgatcg tggctggctc gaagatacct gcaagaatgt
2281 cattgcgctg ccattctcca aattgcagtt cgcgcttagc tggataacgc cacggaatga
2341 tgtcgtcgtg cacaacaatg gtgacttcta cagcgcggag aatctcgctc tctccagggg
2401 aagccgaagt ttccaaaagg tcgttgatca aagctcgccg cgttgtttca tcaagcctta
2461 cggtcaccgt aaccagcaaa tcaatatcac tgtgtggctt caggccgcca tccactgcgg
2521 agccgtacaa atgtacggcc agcaacgtcg gttcgagatg gcgctcgatg acgccaacta
2581 cctctgatag ttgagtcgat acttcggcga tcaccgcttc cctcatactc ttcctttttc
2641 aatattattg aagcatttat cagggttatt gtctcatgag cggatacata tttgaatgta
2701 tttagaaaaa taaacaaata gctagctcac tcggtcgcta cgctccgggc gtgagactgc
```

2761 ggcgggcgct gcggacacat acaaagttac ccacagattc cgtggataag caggggacta
2821 acatgtgagg caaaacagca gggccgcgcc ggtggcgttt ttccataggc tccgccctcc
2881 tgccagagtt cacataaaca gacgcttttc cggtgcatct gtgggagccg tgaggctcaa
2941 ccatgaatct gacagtacgg gcgaaacccg acaggactta aagatcccca ccgtttccgg
3001 cgggtcgctc cctcttgcgc tctcctgttc cgaccctgcc gtttaccgga tacctgttcc
3061 gcctttctcc cttacgggaa gtgtggcgct ttctcatagc tcacacactg gtatctcggc
3121 tcggtgtagg tcgttcgctc caagctgggc tgtaagcaag aactccccgt tcagcccgac
3181 tgctgcgcct tatccggtaa ctgttcactt gagtccaacc cggaaaagca cggtaaaacg
3241 ccactggcag cagccattgg taactgggag ttcgcagagg atttgtttag ctaaacacgc
3301 ggttgctctt gaagtgtgcg ccaaagtccg gctacactgg aaggacagat ttggttgctg
3361 tgctctgcga aagccagtta ccacggttaa gcagttcccc aactgactta accttcgatc
3421 aaaccacctc cccaggtggt tttttcgttt acagggcaaa agattacgcg cagaaaaaaa
3481 ggatctcaag aagatccttt gatcatatga catagtgaca acttgacggc tacatcattc
3541 actttttctt cacaaccggc acggaactcg ctcgggctgg ccccggtgca tttttttaaat
3601 acccgcgaga aatagagttg atcgtcaaaa ccaacattgc gaccgacggt ggcgataggc
3661 atccgggtgg tgctcaaaag cagcttcgcc tggctgatac gttggtcctc gcgccagctt
3721 aagacgctaa tccctaactg ctggcggaaa agatgtgaca gacgcgacgg cgacaagcaa
3781 acatgctgtg cgacgctggc gatatcaaaa ttgctgtctg ccaggtgatc gctgatgtac
3841 tgacaagcct cgcgtacccg attatccatc ggtggatgga gcgactcgtt aatcgcttcc
3901 atgcgccgca gtaacaattg ctcaagcaga tttatcgcca gcagctccga atagcgccct
3961 tccccttgcc cggcgttaat gatttgccca aacaggtcgc tgaaatgcgg ctggtgcgct
4021 tcatccgggc gaaagaaccc cgtattggca aatattgacg gccagttaag ccattcatgc
4081 cagtaggcgc gcggacgaaa gtaaacccac tggtgatacc attcgcgagc ctccggatga
4141 cgaccgtagt gatgaatctc tcctggcggg aacagcaaaa tatcacccgg tcggcaaaca
4201 aattctcgtc cctgattttt caccaccccc tgaccgcgaa tggtgagatt gagaatataa
4261 cctttcattc ccagcggtcg gtcgataaaa aaatcgagat aaccgttggc ctcaatcggc
4321 gttaaacccg ccaccagatg ggcattaaac gagtatcccg gcagcagggg atcattttgc
4381 gcttcagcca tacttttcat actcccgcca ttcagagaag aaaccaattg tccatattgc
4441 atcagacatt gccgtcactg cgtcttttac tggctcttct cgctaaccaa accggtaacc
4501 ccgcttatta aaagcattct gtaacaaagc gggaccaaag ccatgacaaa aacgcgtaac
4561 aaaagtgtct ataatcacgg cagaaaagtc cacattgatt atttgcacgg cgtcacactt
4621 tgctatgcca tagcattttt atccataaga ttagcggatc ctacctgacg ctttttatcg
4681 caactctcta ctgtttctcc atacccgttt tttgggctaa caggaggaac tcgagatggt
4741 tcataaatct gacagtgatg aattagctgc attgagggca gaaaatgccc gtctggtctc
4801 attacttgaa gctcatggga ttgaatggcg acgtaaaccg cagactcctg tgcagcgcgt
4861 ttctgtatta tccaccgacg agaaggttgc attatttcgt cggttgtttc gtgggcgtga
4921 tgatgtatgg gcacttagat gggaaagtaa aaccagcggc aaatcagggt actctcctgc
4981 ctgcgctaac gaatggcagg cgagaatatg cggtaaaccc cggataaaat gtggggactg
5041 cgctcaccgt cagttgattc ctgtatccga tctcgtcatc taccaccatc tggccggtac
5101 ccatactgtc gggatgtatc cgttgctgga agatgattcc tgttattttc tggcagttga
5161 ttttgatgaa gctgaatggc agaaggatgc atccgcattc atgcgatcct gtgatgagct
5221 gggtgtacct gctgcgctgg aaatatcccg ttcacgtcag ggggcgcatg tctggatatt
5281 ttttgcctca cgagtttcgg cccgcgaagc tcgccgtctg ggactgcta ttattagcta
5341 tacgtgtagt cggacccgac agctgcgatt ggggtcttat gaccgattat ttcctaatca
5401 ggatactatg ccaaaagggg gatttggtaa tctcatagcg ttacctctgc aaaaaagacc
5461 gcgtgaattg gggggaagcg tttttgttga tatgaatctc cagccttatc ctgatcagtg
5521 ggcttttctt gtatcggtga ccccgatgaa tgtgcaggat attgaaccga cgatattacg
5581 ggctacaggg agtatccatc ctctggatgt gaattttatc aacgaagaag acctgggtac
5641 gccgtgggaa gagaaaaaat catcaggaaa cagactgaat atttctattg cagaaccgct
5701 gaaaatcacg ctggcaaacc agatctattt cgaaaaagcg caattacctc aggtgctgat
5761 taaccgactt attcggctgg cagcatttcc gaaccctgag ttttataagg ctcaggcaat
5821 gcgtatgtca gtctggaata agccccgtgt tataggttgt gcggagaatt acccgcaaca
5881 cattgcgttg ccccggggat gtctggacag cgtattatct ttccttaggg acaacaatat
5941 tgctgcagaa ttaatcgata aacgatttgc cgggacggaa tgtaatgccg tttttatggg
6001 aaacctcaga gcggagcagg aagaggccgt ttcggcatta ctccgttatg acactggtgt
6061 gctttgtgcg ccaacggctt ttggtaagac agttaccgca gcggcagtga ttgccaagcg
6121 gaaagtgaat acactgatac tggtacaccg gactgaattg ctgaagcagt ggcaggagcg
6181 tctcgcggtg tttctgcagg ccggtgacag tattggtatt atcgggggag gtaaacataa
6241 accctgtggc aatattgata ttgcggtggt gcagtccata tccagacaag gagaagttga

```
6301 acctctggtc aggaattatg ggcaaatcat tgtggatgag tgccatcata ttggcgcggt
6361 ttcattttct gcgattctga aggaaacgaa tgccagatat ctgcttggcc tgacggcaac
6421 accaatccga cgggatggtc tgcatcccat tatttttatg tactgtggtg ccattcgcca
6481 tacagcggtc cgcccgaagg aaagcccaca taatctggag gtactgatcc gttcccgttt
6541 tacatctggt catttaccat cggatgcgag aatccaggat attttcagag aaattgctct
6601 ggatcatgac agaacggtgg cgatagctga agaagccatg aaagctttcg ggcaggggcg
6661 aaaagttctg gtactgactg aacgtacaga tcatctggat gagatagcat cagtgatgaa
6721 ttcactgaaa ttgtctccct ttattctcca tggtcgacta tcgaaaaaaa agcgtgcgat
6781 gctgatatcc gggctgaatg ctcttcctcc cgattctcct cgaattttgt tgtcaacagg
6841 cagacttatt ggtgagggat ttgaccaccc tccgctggat acgctgattc ttgccatgcc
6901 tgtgtcatgg aaagggacat tacagcagta tgcagggcgt cttcacagag agcataccgg
6961 taaaagcgat gtcaggatca ttgattttgt ggataccgcg tatcctgtgc tgctcagaat
7021 gtgggataaa cgtcagcggg gttataaagc gatggggtac aggattatag ctgacggcga
7081 tgaatcagtg atttaagggg gattcacctt tatgttgata agaaataaaa gaaaatgcca
7141 ataggatatc ggcattttct tttgcgttt
//
```

Sequence between *kanR* and origin of replication to make *kanR* of pYK-1N5 inducible with IPTG

```
   1 ccccagagtc ccgctcagaa gaactcgtca agaaggcgat agaaggcgat gcgctgcgaa
  61 tcgggagcgg cgataccgta aagcacgagg aagcggtcag cccattcgcc gccaagctct
 121 tcagcaatat cacgggtagc caacgctatg tcctgatagc ggtccgccac acccagccgg
 181 ccacagtcga tgaatccaga aaagcggcca ttttccacca tgatattcgg caagcaggca
 241 tcgccatggg tcacgacgag atcctcgccg tcgggcatgc gcgccttgag cctggcgaac
 301 agttcggctg gcgcgagccc ctgatgctct tcgtccagat catcctgatc gacaagaccg
 361 gcttccatcc gagtacgtgc tcgctcgatg cgatgtttcg cttggtggtc gaatgggcag
 421 gtagccggat caagcgtatg cagccgccgc attgcatcag ccatgatgga tactttctcg
 481 gcaggagcaa ggtgagatga caggagatcc tgccccggca cttcgcccaa tagcagccag
 541 tcccttcccg cttcagtgcc aacgtcgagc acagctgcgc aaggaacgcc cgtcgtggcc
 601 agccacgata gccgcgctgc ctcgtcctgc agttcattca gggcaccgga caggtcggtc
 661 ttgacaaaaa gaaccgggcg cccctgcgct gacagccgga acacggcggc atcagagcag
 721 ccgattgtct gttgtgccca gtcatagccg aatagcctct ccacccaagc ggccggagaa
 781 cctgcgtgca atccatcttg ttcaatcatc atctagtatt tcccctcttt ctctagtatt
 841 aaacaaaatt atttgtagag gctgtttcgt cctcacggac tcatcagacc ggaaagcaca
 901 tccggtgaca gctaattgtg agcgctcaca attccacaca ttatacgagc cgatgattaa
 961 ttgtcaacac tcggtaccaa attccagaaa agaggcctcc cgaaaggggg gccttttttc
1021 gttttggtcc aatggcggcg cgccatcgaa tggcgcaaaa cctttcgcgg tatggcatga
1081 tagcgcccgg aagagagtca attcagggtg gtgaatatga aaccagtaac gttatacgat
1141 gtcgcagagt atgccggtgt ctcttatatg accgtttccc gcgtggtgaa ccaggccagc
1201 cacgtttctg cgaaaacgcg ggaaaaagtg gaagcggcga tggtggagct gaattacatt
1261 cccaaccgcg tggcacaaca actggcgggc aaacagtcgt tgctgattgg cgttgccacc
1321 tccagtctgg ccctgcacgc gccgtcgcaa attgtcgcgg cgattaaatc tcgcgccgat
1381 caactgggtg ccagcgtggt ggtgtcgatg gtagaacgaa gcggcgtcga agcctgtaaa
1441 gcggcggtgc acaatcttct cgcgcaacgc gtcagtgggc tgatcattaa ctatccgctg
1501 gatgaccagg atgccattgc tgtggaagct gcctgcacta atgttccggc gttatttctt
1561 gatgtctctg accagacacc catcaacagt attattttct cccatgagga cggtacgcga
1621 ctgggcgtgg agcatctggt cgcattgggt caccagcaaa tcgcgctgtt agcgggccca
1681 ttaagttctg tctcggcgcg tctgcgtctg gctggctggc ataaatatct cactcgcaat
1741 caaattcagc cgatagcgga acgggaaggc gactggagtg ccatgtccgg ttttcaacaa
1801 accatgcaaa tgctgaatga gggcatcgtt cccactgcga tgctggttgc caacgatcag
1861 atggcgctgg gcgcaatgcg cgccattacc gagtccgggc tgcgcgttgg tgcggatatc
1921 tcggtagtgg gatacgacga taccgaagat agctcatgtt atatcccgcc gttaaccacc
1981 atcaaacagg attttcgcct gctggggcaa accagcgtgg accgcttgct gcaactctct
2041 cagggccagg cggtgaaggg caatcagctg ttgccagtct cactggtgaa aagaaaaacc
2101 accctggcgc ccaatacgca aaccgcctct ccccgcgcgt tggccgattc attaatgcag
2161 ctggcacgac aggtttcccg actggaaagc gggcagtgat aaggatccta attggtaacg
2221 aatcagacaa ttgacggctc gagggagtag catagggttt gcagaatccc tgcttcgtcc
2281 atttgacagg cacattatgc atcgatgata agctgtcaaa catgagcaga tcctctacgc
2341 cggacgcatc gtggccggca tcaccggcgc cacaggtgcg gttgctggcg cctatatcgc
2401 cgacatcacc gatggggaag atcgggctcg ccacttcggg ctcatgagca aatattttat
```

```
     2461 ctgaggtgct tcctcgctca ctgacgcgga acccctattt gtttattttt ctaaatacat
     2521 tcaaatatgt atccgctcat gagacaataa ccctgataaa tgcttcaata atattgaaaa
     2581 aggaagagta tgagtattca acatttccgt gtcgccctta ttcccttttt tgcggcattt
     2641 tgccttcctg tttttgctca cccagaaacg ctggtgaaag taaaagatgc tgaagatcag
     2701 ttgggtgcac gagtgggtta catcgaactg gatctcaaca gcggtaagat ccttgagagt
     2761 tttcgccccg aagaacgttt tccaatgatg agcactttta aagttctgct atgtggcgcg
     2821 gtattatccc gtattgacgc cgggcaagag caactcggtc gccgcataca ctattctcag
     2881 aatgacttgg ttgagtactc accagtcaca gaaaagcatc ttacggatgg catgacagta
     2941 agagaattat gcagtgctgc cataaccatg agtgataaca ctgcggccaa cttacttctg
     3001 acaacgatcg gaggaccgaa ggagctaacc gcttttttgc acaacatggg ggatcatgta
     3061 actcgccttg atcgttggga accggagctg aatgaagcca taccaaacga cgagcgtgac
     3121 accacgatgc ctgtagcaat ggcaacaacg ttgcgcaaac tattaactgg cgaactactt
     3181 actctagctt cccggcaaca attaatagac tggatggagg cggataaagt tgcaggacca
     3241 cttctgcgct cggcccttcc ggctggctgg tttattgctg ataaatctgg agccggtgag
     3301 cgtgggtctc gcggtatcat tgcagcactg gggccagatg gtaagccctc ccgtatcgta
     3361 gttatctaca cgacggggag tcaggcaact atggatgaac gaaatagaca gatcgctgag
     3421 ataggtgcct cactgattaa gcattggtaa ccaggcatca aataaaacga aaggctcagt
     3481 cgaaagactg ggcctttcgt tttatctgtt gtttgtcggt gaactgtaag acgggtaagc
     3541 ctgttgatga taccgctgcc ttactgggtg cattagccag tctgaatgac ctgtcacggg
     3601 ataatccgaa gtggtcagac tggaaaatca gagggcagga actgcagaac agcaaaaagt
     3661 cagatagcac cacatagcag acccgccata aaacgccctg agaagcccgt gacgggcttt
     3721 tcttgtatta tgggtagttt ccttgcatga atccataaaa ggcgcctgta gtgccattta
     3781 cccccattca ctgccagagc cgtgagcgca gcgaactgaa tgtcacgaaa aagacagcga
     3841 ctcaggtgcc tgatggtcgg agacaaaagg aatattcagc gatttgcccg agcttgcgag
     3901 ggtgctactt aagcctttag ggttttaagg tctgttttgt agaggagcaa acagcgtttg
     3961 cgacatcctt ttgtaatact gcggaactga ctaaagtagt gagttataca cagggctggg
     4021 atctattctt tttatctttt tttattcttt ctttattcta taaattataa ccacttgaat
     4081 ataaacaaaa aaaacacaca aaggtctagc ggaatttaca gagggtctag cagaatttac
     4141 aagttttcca gcaaaggtct agcagaattt acagataccc acaactcaaa ggaaaaggac
     4201 tagtaattat cattgactag cccatctcaa ttggtatagt gattaaaatc acctagacca
     4261 attgagatgt atgtctgaat tagttgtttt caaagcaaat gaactagcga ttagtcgcta
     4321 tgacttaacg gagcatgaaa ccaagctaat tttatgctgt gtggcactac tcaaccccac
     4381 gattgaaaac cctacaagga aagaacggac ggtatcgttc acttataacc aatacgctca
     4441 gatgatgaac atcagtaggg aaaatgctta tggtgtatta gctaaagcaa ccagagagct
     4501 gatgacgaga actgtggaaa tcaggaatcc tttggttaaa ggctttgaga ttttccagtg
     4561 gacaaactat gccaagttct caagcgaaaa attagaatta gtttttagtg aagagatatt
     4621 gccttatctt ttccagttaa aaaaattcat aaaatataat ctggaacatg ttaagtcttt
     4681 tgaaaacaaa tactctatga ggatttatga gtggttatta aaagaactaa cacaaaagaa
     4741 aactcacaag gcaaatatag agattagcct tgatgaattt aagttcatgt taatgcttga
     4801 aaataactac catgagttta aaaggcttaa ccaatgggtt ttgaaaccaa taagtaaaga
     4861 tttaaacact tacagcaata tgaaattggt ggttgataag cgaggccgcc cgactgatac
     4921 gttgattttc caagttgaac tagatagaca aatggatctc gtaaccgaac ttgagaacaa
     4981 ccagataaaa atgaatggtg acaaaatacc aacaaccatt acatcagatt cctacctaca
     5041 taacggacta agaaaaacac tacacgatgc tttaactgca aaaattcagc tcaccagttt
     5101 tgaggcaaaa tttttgagtg acatgcaaag taagcatgat ctcaatggtt cgttctcatg
     5161 gctcacgcaa aaacaacgaa ccacactaga gaacatactg gctaaatacg gaaggatctg
     5221 aggttcttat ggctcttgta tctatcagtg aagcatcaag actaac
//
```
